# A genome-wide screen for ER autophagy highlights key roles of mitochondrial metabolism and ER-resident UFMylation

**DOI:** 10.1101/561001

**Authors:** Jin Rui Liang, Emily Lingeman, Thao Luong, Saba Ahmed, Truc Nguyen, James Olzmann, Jacob E. Corn

**Affiliations:** Innovative Genomics Institute, University of California, Berkeley, Berkeley, California 94720, USA; Department of Molecular and Cell Biology, University of California, Berkeley, Berkeley, California 94720, USA; Department of Nutritional Sciences and Toxicology, University of California, Berkeley, Berkeley, California 94720, USA

## Abstract

Selective degradation of organelles via autophagy is critical for cellular differentiation, homeostasis, and organismal health. Autophagy of the ER (ER-phagy) is implicated in human neuropathy but is poorly understood beyond a few specialized autophagosomal receptors and remodelers. Using an ER-phagy reporter and genome-wide CRISPRi screening, we identified 200 high-confidence factors involved in human ER-phagy. We mechanistically investigated two pathways unexpectedly required for ER-phagy. First, reduced mitochondrial metabolism represses ER-phagy, which reverses the logic of general autophagy. Mitochondrial crosstalk with ER-phagy bypasses the energy sensor AMPK, instead directly impacting ULK1. Second, ER-localized UFMylation is required for ER-phagy that represses the unfolded protein response. The UFL1 ligase is brought to the ER surface by DDRGK1, analogous to PINK1-Parkin regulation during mitophagy. Our data provide insight into the unique cellular logic of ER-phagy, reveal parallels between organelle autophagies, and provide an entry point to the relatively unexplored process of degrading the ER network.

## Introduction

Macroautophagy (herein referred to as autophagy) mediates the delivery of cellular cargo to the lysosome for degradation. Once thought to be a non-specific process, it has become clear that autophagy is complexly regulated and induced by various stresses to remove damaged or excessive cellular components. Targeted removal of entire organelles by autophagy is necessary for cellular homeostasis, and during selective autophagy of mitochondria (mitophagy), the surface proteins of damaged mitochondria are marked by phosphorylation and ubiquitylation to recruit autophagic machinery (Nguyen et al., 2016; Youle and Narendra, 2011). Dysregulation of selective organelle autophagy negatively impacts cellular fitness and is linked to degenerative diseases, particularly in non-regenerative cell types such as neurons. Mutations of key mitophagy genes, such as PINK1 and Parkin, are strongly associated with disorders such as Parkinson’s disease (Deas et al., 2011; Dodson and Guo, 2007; Geisler et al., 2010; Pickrell and Youle, 2015; Pilsl and Winklhofer, 2012).

The endoplasmic reticulum (ER) plays a critical role in numerous cellular functions, such as the folding, modification, and transport of secretory proteins; the storage of calcium; and the biosynthesis of lipids (Schwarz and Blower, 2016). The ER is tightly regulated by multiple quality control mechanisms, such as ER-associated degradation (ERAD) and ER to Lysosome Associated Degradation (ERLAD) (Fregno et al., 2018; Ruggiano et al., 2014). The ER-autophagy (ER-phagy) pathway intersects with the selective autophagy machinery to send portions of the ER for wholesale lysosomal degradation. While ER-phagy has long been observed in yeast (Hamasaki et al., 2005), it has only recently been described in mammalian cells (Khaminets et al., 2015).

During ER-phagy at least two ER surface proteins, FAM134B and RTN3L, act as specific receptors through LC3-interacting regions (LIR) that recruit autophagy machinery (Grumati et al., 2017; Khaminets et al., 2015). ER expansion can also be reversed via ER-phagy that is mediated by the SEC62 and CCPG1 LIR-containing ER-phagy receptors (Fumagalli et al., 2016; Smith et al., 2018). The reticular ER network is remodeled for delivery to the lysosome by Atlastin GTPases that are also involved in normal ER morphology (Liang et al., 2018; Rismanchi et al., 2008; Wang et al., 2016; Zhao et al., 2016). Failure to execute ER-phagy through mutations in known ER-phagy proteins causes neuropathy in mice, and cross-referencing with ClinVar reveals that mutations in human ER-phagy proteins are linked to hereditary neuropathies and paraplegia (Guelly et al., 2011; Khaminets et al., 2015; Kurth et al., 2009; Liang et al., 2018). But beyond the few receptors and remodelers most proximal to autophagosomal function, relatively little is known about the signals that regulate ER-phagy.

We performed a genome-wide CRISPR interference (CRISPRi) reporter-based screen to identify new players in ER-phagy, identifying both activators and inhibitors in a variety of cellular compartments. We mechanistically interrogated two pathways that positively regulate ER-phagy in unexpected ways: (1) mitochondrial oxidative phosphorylation (OXPHOS) and (2) ER-resident UFMylation. While inhibition of OXPHOS reduces cellular energy levels and stimulates general autophagy, genetic or chemical inhibition of OXPHOS instead represses ER-phagy. Mitochondrial regulation of ER-phagy bypasses the canonical energy-sensing AMP-dependent protein kinase (AMPK), and instead directly modulates ULK1 levels during starvation. We furthermore found that UFMylation, a post-translational modification similar to ubiquitination, is required for ER-phagy. The protein DDRGK1 recruits the UFMylation machinery to the ER surface in a striking parallel to the mitophagic recruitment of Parkin by PINK1. Interfering with DDRGK1-dependent UFMylation inhibits ER-phagy, leading to the accumulation of misfolded proteins and inducing the unfolded protein response (UPR) via IRE1α signaling. Overall, our data provide a detailed map of ER-phagy regulators and highlight how organelle crosstalk and ER-resident factors mediate this emerging process of quality control.

## Results

### A genome-wide flow cytometry CRISPRi screen for ER-phagy

To develop a genome-wide screen for ER-phagy, we employed the previously-developed ER-Autophagy Tandem Reporter (EATR) system (Fig 1A) (Liang et al., 2018). In HCT116 colon cancer cells that stably express the EATR construct, we found that inducing general autophagy via inactivation of the mTOR complex did not induce ER-phagy (Fig S1A-B). Similarly, ER stress-inducing drugs that induce the unfolded protein response (UPR) did not cause ER-phagy. Only prolonged starvation (16 hours) using Earl’s Buffered Saline Solution (EBSS) robustly induced ER-phagy.

**Figure 1.**
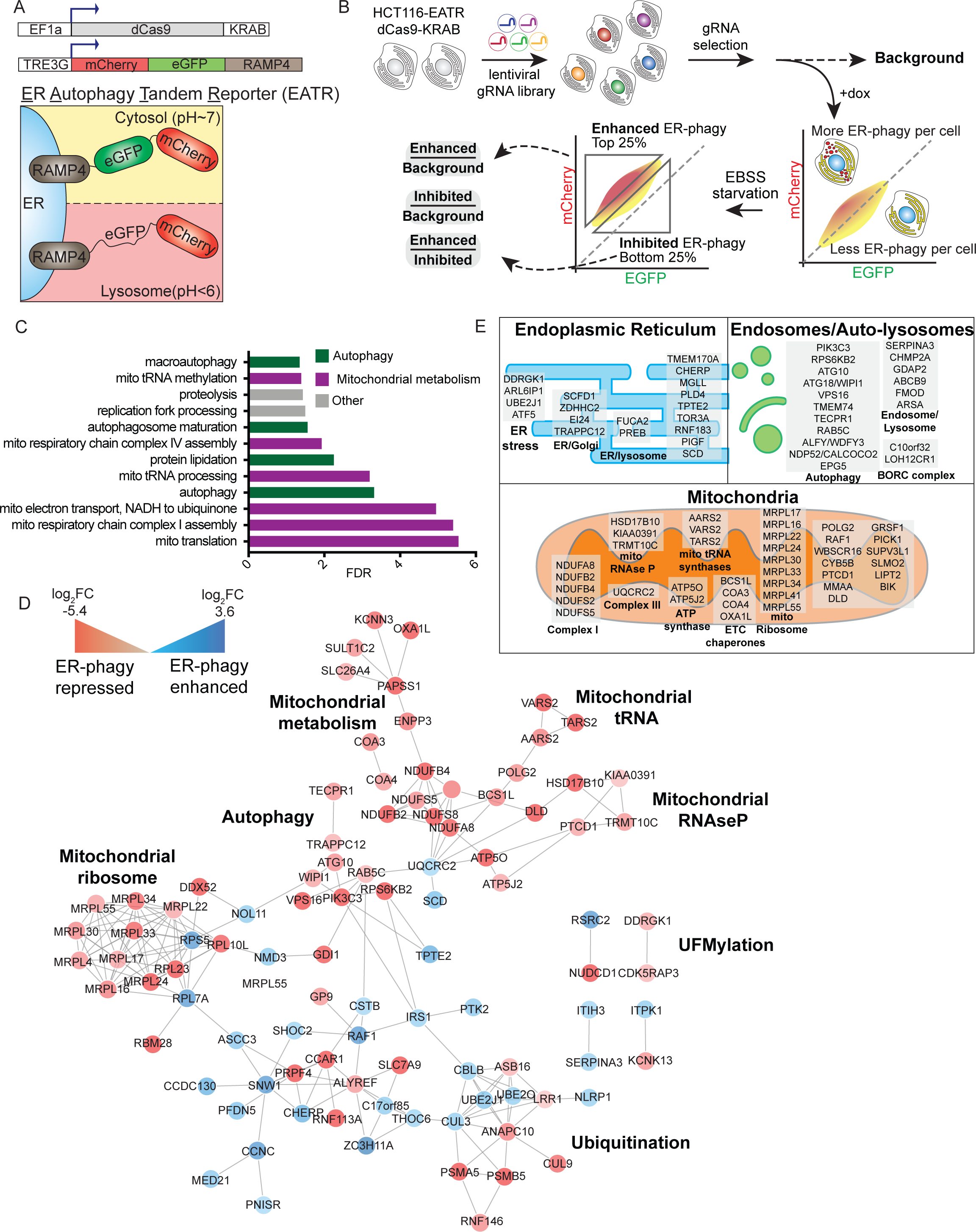
Unbiased identification of human ER-phagy regulators by genome-wide CRISPRi screening. A. Schematic of the <u>E</u>R <u>A</u>utophagy <u>T</u>andem <u>R</u>eporter (EATR) and CRISPR inhibition (CRISPRi) system used for screening. HCT116 cells stably express a doxycycline-inducible EATR construct that consists of mCherry and eGFP fused to ER localized RAMP4. Cells also stably express dCas9-KRAB for gRNA-targeted transcriptional repression of targeted genes.
B. FACS screening strategy for identifying human genes whose knockdown enhances or inhibits ER-phagy. HCT116 cells stably expressing dCas9-KRAB and dox-inducible EATR were transduced with a genome-wide lentiviral CRISPRi gRNA library. After selection for gRNA expression and removal of essential genes, doxycycline was added for 24 hr to express EATR. Cells were then treated with EBSS starvation media for 16 hr to induce ER-phagy. The top and bottom quartiles of cells correspond enhanced or inhibited ER-phagy in response to CRISPRi knockdown. Cells were sorted and processed for next generation sequencing to identify gRNA representation in each sort bin.
C. Gene ontology analysis identifies autophagy and mitochondrial metabolism as major signatures of ER-phagy. High-confidence ER-phagy genes were defined as having opposite phenotypes in the enhanced and inhibited sort gates and gene-level p < 0.01. Ontologies with Benjamini-Hochberg FDR < 0.05 are shown.
D. Genes involved in ER-phagy form a physical interaction network. For clarity, only interactions between two or more high-confidence hits are shown. Red circles represent genes whose knockdown represses ER-phagy and blue circles represent genes whose knockdown enhances ER-phagy. The shades of red/blue correspond to the log2-fold-change associated with each gene.
E. Subcellular classification of high-confidence ER-phagy genes highlights roles in the ER, auto-lysosomes, and mitochondria.

Using EBSS as an ER-phagy stimulus, we coupled EATR-based FACS screening with genome-wide CRISPR transcriptional inhibition (CRISPRi) to identify novel pathways involved in ER-phagy (Fig 1A-B) (Gilbert et al., 2014; Horlbeck et al., 2016; Liang et al., 2018). Since autophagy is influenced by the availability of cellular energy, we reasoned that complete knockout of ER-phagy regulators via CRISPR cutting could be detrimental to cells and mask interesting players. The variability in sgRNA efficiencies of CRISPRi leads to different knockdown efficiencies, allowing for allelic series and residual function of essential genes involved in cellular energy regulation (Horlbeck et al., 2016).

As a proof-of-concept, we first assessed the suitability of EATR for CRISPRi screening by conducting a pilot screen with a custom CRISPRi library targeting known autophagy genes (Table S1). We used EATR-based FACS to isolate the top 25% of cells with the most ER-phagy (‘enhanced’ sort gate), and the bottom 25% of cells with the least ER-phagy (‘inhibited’ sort gate) (Fig S1C). This pilot screen successfully identified gRNAs targeting core autophagy genes as required for ER-phagy, and correctly assigned their role in promoting ER-phagy such that knockdown of autophagy components was enriched in the ‘inhibited’ gate (Fig S1D) and depleted in the ‘enhanced’ sort gate (Fig S1E).

We scaled up to perform an unbiased, genome-wide CRISPRi-v2 screen (Gilbert et al., 2014; Horlbeck et al., 2016) for ER-phagy regulators using EATR-FACS. From the gene-level statistics, we defined a high-confidence list of ER-phagy genes by first performing a cutoff at p < 0.01, and then requiring that true hits have opposite phenotypes in the “enhanced” and “inhibited” sort gates relative to the unsorted background. For example, gRNAs that knockdown a *bona fide* gene required for ER-phagy should be depleted in a population undergoing more ER-phagy but enriched in one undergoing less ER-phagy. We quantified involvement in ER-phagy by ratiometrically comparing gRNA distributions in the enhanced gate to those in the inhibited gate (Fig 1B), so that a positive log_2_-fold-change indicates a gene whose knockdown increases ER-phagy and a negative log_2_-fold-change indicates a gene whose knockdown inhibits ER-phagy. The resulting high-confidence hit list includes 200 genes involved in ER-phagy, with gene-level log_2_-fold-change phenotypes ranging from 3.62 to −5.38 (Table S2).

As expected, multiple stages of general autophagy and membrane trafficking were prominent hits from the genome-wide library (Fig S1F). Individual stable knockdown of these autophagy-related factors and testing using EATR, an m<u>C</u>herry-<u>C</u>leaved <u>E</u>R-phagy <u>R</u>eporter (CCER) western blot assay, and other measures of autophagy verified their requirement for ER-phagy (Fig S1G-I). These data indicate that the EATR assay and hence the genome-wide screen reports on ER-phagy pathways rather than ERAD or ERLAD, since the latter do not depend on autophagic components (Fregno et al., 2018; Ruggiano et al., 2014). We noticed that some of the established ER-phagy receptors such as FAM134B were not gene-level hits, and examining the performance of individual gRNAs by qRT-PCR, we found that none of the CRISPRi gRNAs in the genome-wide library successfully knocks down FAM134B (Fig S1J). This highlights a tradeoff in current CRISPRi screening technology, where allelic series enable interrogation of otherwise essential genes but may introduce false negatives. Failure to observe a gene in the ER-phagy screen dataset therefore does not imply that it plays no role in ER-phagy, but positive membership in the high-confidence set of 200 genes does indicate a role in ER-phagy.

We used unbiased GO term analysis of the high-confidence gene-level hit list to determine the cellular functions and pathways involved in ER autophagy (Fig 1C) (Huang et al., 2009). As expected for ER-phagy, GO terms related to autophagy were significantly enriched. However, we were surprised to find that multiple aspects of mitochondrial metabolism were even more prominent. We cross-referenced the high-confidence genetic hits against physical interaction databases to create a putative physical network of ER-phagy (Chatr-Aryamontri et al., 2017; Szklarczyk et al., 2015) (Fig 1D). This network falls into several major classes: autophagic execution, such as ATG10, (Phillips et al., 2008; Wartosch et al., 2015; Yuan et al., 2013); ubiquitination, such as the ER-localized UBE2J1 ubiquitin-conjugating enzyme involved in recovery from ER stress (Elangovan et al., 2017); mitochondrial metabolism, including nuclear-encoded OXPHOS genes, OXPHOS chaperones, and components of the mitochondrial ribosome required for translation of mitochondrially-encoded OXPHOS genes; and post-translational modification by the ubiquitin-like protein UFM1, including CDK5RAP3 and DDRGK1 (Cai et al., 2015; Wei and Xu, 2016; Wu et al., 2010). Finally, we manually annotated all 200 high-confidence hits based on their subcellular localization (Binder et al., 2014). The localization analysis subdivided factors involved in ER-phagy into those associated with the lysosome/endosome, ER-associated factors, and nuclear-encoded mitochondrial proteins (Fig 1E).

### Mitochondrial oxidative phosphorylation promotes ER-phagy by regulating levels of ULK1

We were surprised to find that the largest set of genes required for ER-phagy are involved in OXPHOS (Fig 2A), since cross-talk between ER-phagy and mitochondrial processes has not yet been described. While interference with mitochondrial energy metabolism induces general cytoplasmic autophagy, loss of mitochondrial factors instead repressed ER-phagy. Mitochondrial factors required for ER-phagy are directly associated with multiple aspects of the electron transport chain (ETC) and OXPHOS: Complex I (NDUFA8, NDUFB2, NDUFB4, NDUFS2, NDUFS5), Complex III (UQCRC2), and the ATP synthase/Complex V (ATP5O and ATP5J) (Fig S2A). We also found a large number of factors indirectly required for OXPHOS either through ETC maturation or the synthesis of mitochondrially-encoded ETC components (Taanman, 1999): mitochondrial chaperones (BCS1L, COA3, COA4, and OXA1L), mitochondrial ribosome subunits (MRPL17, MRPL16, MRPL22, MRPL24, MRPL30, MRPL33, MRPL34, MRPL41, MRPL55), mitochondrial tRNAs (AARS2, VARS2, TARS2), mitochondrial tRNA maturation (PTCD1 and mitochondrial RNAse P KIAA0391, TRMT10C, and HSD17B10). To further interrogate the link between mitochondrial metabolism and ER-phagy, we focused on mechanistic characterization of three factors that are involved in different parts of OXPHOS: NDUFB2, NDUFB4, and ATP5O.

**Figure 2.**
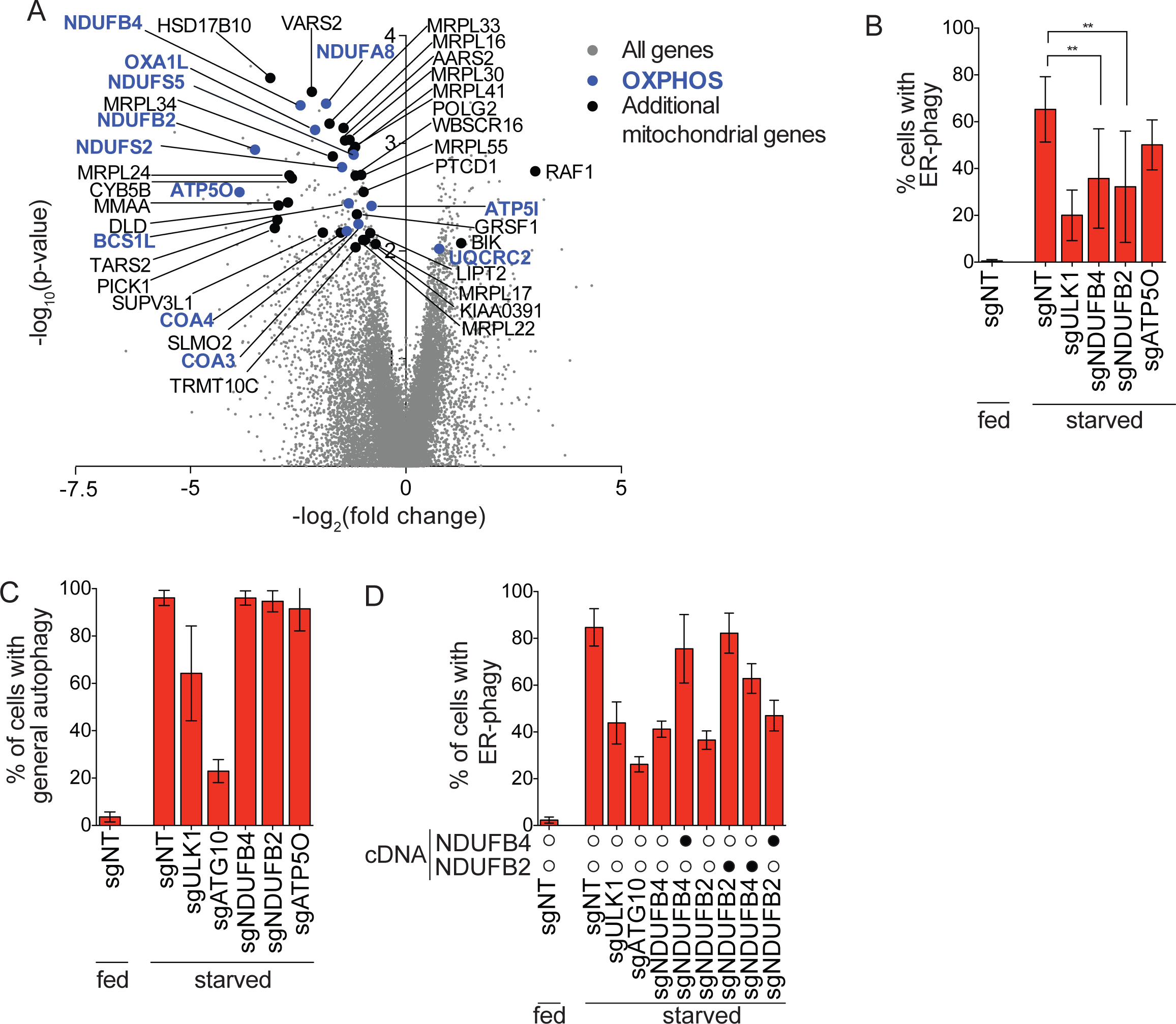
Intact OXPHOS promotes ER-phagy. A. Genes that are components of the OXPHOS pathway were top hits in screen (highlighted in blue). Additional mitochondria-related genes are indicated in black and all other targeting sgRNAs are indicated in grey.
B. Knockdown of NDUFB4 and NDUFB2 significantly inhibit ER-phagy. HCT116 CRISPRi EATR cells were transduced with sgRNAs targeting ULK1, NDUFB4, NDUFB2, or ATP5O and starved for 16hr before FACS measurement for ER-phagy. Data presented as mean ± SD of eight biological replicates. P value indicates two-tailed paired t-test (**, P < 0.01).
C. General autophagy proceeds in cells where NDUFB2, NDUFB4, or ATP4O are knocked down. HCT116 CRISPRi cells expressing eGFP-mCherry-LC3B were transduced with sgRNAs targeting either ULK1, ATG10, NDUFB4, NDUFB2, or ATP5O. Cells were starved for 16hr before FACS measurement for general autophagy. Data represents mean ± SD of two biological replicates.
D. Re-expression of NDUFB4 and NDUFB2 rescues ER-phagy. HCT116 CRISPRi EATR cells were transduced with NDUFB4 or NDUFB2 cDNA constructs, and then transduced with sgRNAs targeting ULK1, ATG10, NDUFB4, NDUFB2. Cells were starved for 16hr before FACS measurement for ER-phagy. Data presented as mean ± SD of three biological replicates.

Individual, stable knockdown of OXPHOS components quantitatively inhibited starvation-induced ER-phagy in multiple assays in a manner that paralleled knockdown efficiency (Fig 2B, S2B). However knockdown of NDUFB2, NDUFB4, or ATP5O did not inhibit starvation-induced general autophagy (Fig 2C, Fig S2C). Re-expressing the appropriate OXPHOS cDNA rescued ER-phagy, while cross-expressing a non-cognate OXPHOS cDNA had no effect, indicating that each knockdown was specific and on target (Fig 2D, Fig S2D-F). We further explored the necessity of functional OXPHOS for normal ER-phagy using chemical genetics (Fig 3A). Rotenone is a known inhibitor of both general autophagy and Complex I (Mader et al., 2012), and reduced both general autophagy and ER-phagy by multiple assays (Fig 3B – E, Fig S2G). By contrast, inhibiting Complex III with antimycin A or ATP synthase with oligomycin A increased general autophagy but reduced ER-phagy (Fig 3B – E, Fig S2G – H). Using Cell-Titer Glo to measure ATP levels (Fig. S2I) and Seahorse to measure oxygen consumption (Fig. S2J), we confirmed that knockdown of NDUFB2, NDUFB4, or ATP5O reduced cellular energy levels.

**Figure 3.**
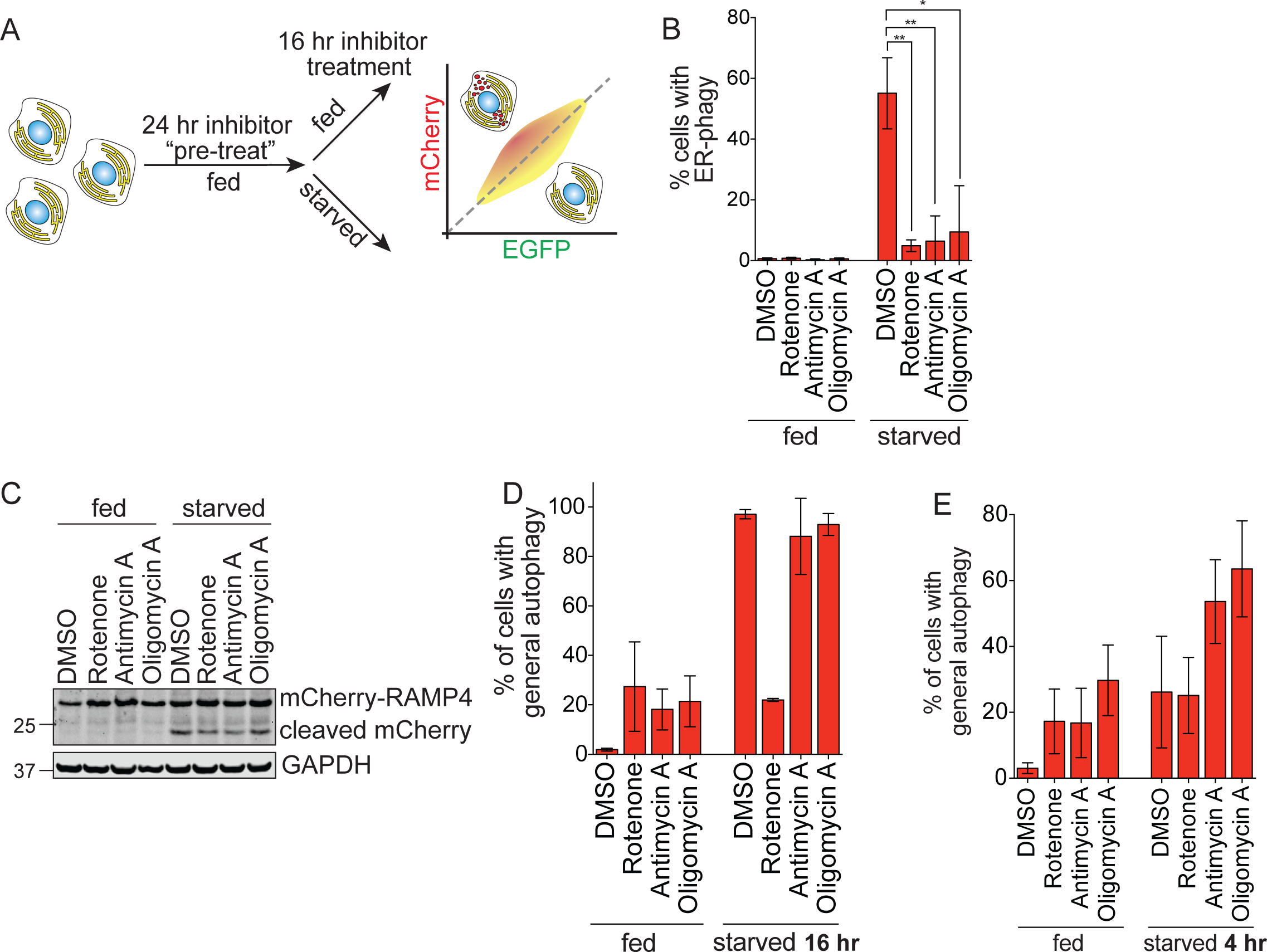
Electron transport chain inhibitors impair ER-phagy. A. Schematic of experimental setup for chemical genetic inhibition of electron transport chain function in ER-phagy. HCT116 CRISPRi cells were treated with each inhibitor for 24 hours in complete media, then maintained in inhibitor for another 16 hours in fed or EBSS starvation media. Cells were then harvested for FACS to measure autophagy.
B. OXPHOS inhibitors disrupt ER-phagy as measured by flow cytometry. HCT116 EATR cells were treated with small molecule inhibitors of rotenone, antimycin A, or oligomycin A, and starved for 16 hours before FACS measurement of ER-phagy. Data presented as mean ± SD of three biological replicates. P value indicates two-tailed unpaired t-test (*, P < 0.05, **, P < 0.01).
C. OXPHOS inhibitors disrupt ER-phagy as measured by Western blot. HCT116 CRISPRi CCER cells were treated with rotenone, antimycin A, and oligomycin A and starved for 16hr. Cells were lysed for Western blotting to measure mCherry-RAMP4 cleavage.
D. Rotenone, a known autophagy inhibitor, inhibits general autophagy, while antimycin A and oligomycin A have no effect at time points where ER-phagy is disrupted. HCT116 CRISPRi cells expressing eGFP-mCherry-LC3B were treated with rotenone, antimycin A, or oligomycin A and starved for 16hr before FACS measurement for general autophagy. Data represents mean ± SD of three biological replicates.
E. Antimycin A and oligomycin A promote general autophagy at short time points. HCT116 CRISPRi cells expressing eGFP-mCherry-LC3B were treated with rotenone, antimycin A, or oligomycin A and starved for 4hr before FACS measurement for general autophagy. Data represents mean ± SD of three biological replicates.

Depolarized mitochondria can be degraded via mitophagy, and dysregulation of OXPHOS might lead to mitochondrial depolarization by reducing the proton gradient across the mitochondrial inner membrane. Using Mitotracker, we found that OXPHOS conditions that repressed ER-phagy did not grossly alter mitochondrial abundance or membrane potential (Fig S3A). Consistent with prior reports, starvation actually increased the amount of Mitotracker accumulation (Johnson et al., 2014; Xiao et al., 2016). We furthermore found no evidence of increased mitophagy during knockdown of OXPHOS components to repress ER-phagy, either with or without stable over-expression of Parkin (Fig S3B – F). We therefore disfavor a model in which OXPHOS depletion inhibits ER-phagy through potent upregulation of mitophagy.

The inhibition of ER-phagy by interfering with OXPHOS is counter-intuitive, since a reduction in energy levels is generally accepted to induce bulk cytoplasmic autophagy via 5’ AMP-activated Protein Kinase (AMPK) signaling. Reductions in cellular ATP levels activate AMPK, which has also recently been implicated in the regulation of mitophagy (Egan et al., 2011; Kim et al., 2011; Toyama et al., 2016). We therefore asked whether the reduced energy levels caused by repression of OXPHOS leads to AMPK activity that somehow stimulates general autophagy while simultaneously repressing ER-phagy. However, we were surprised to find that AMPKα CRISPR-Cas9 knockout cells still mount a robust ER-phagy response, and this response is unaffected by stable re-expression of constitutively active or kinase dead AMPKα (Fig. 4A-B, Fig S3G).

**Figure 4.**
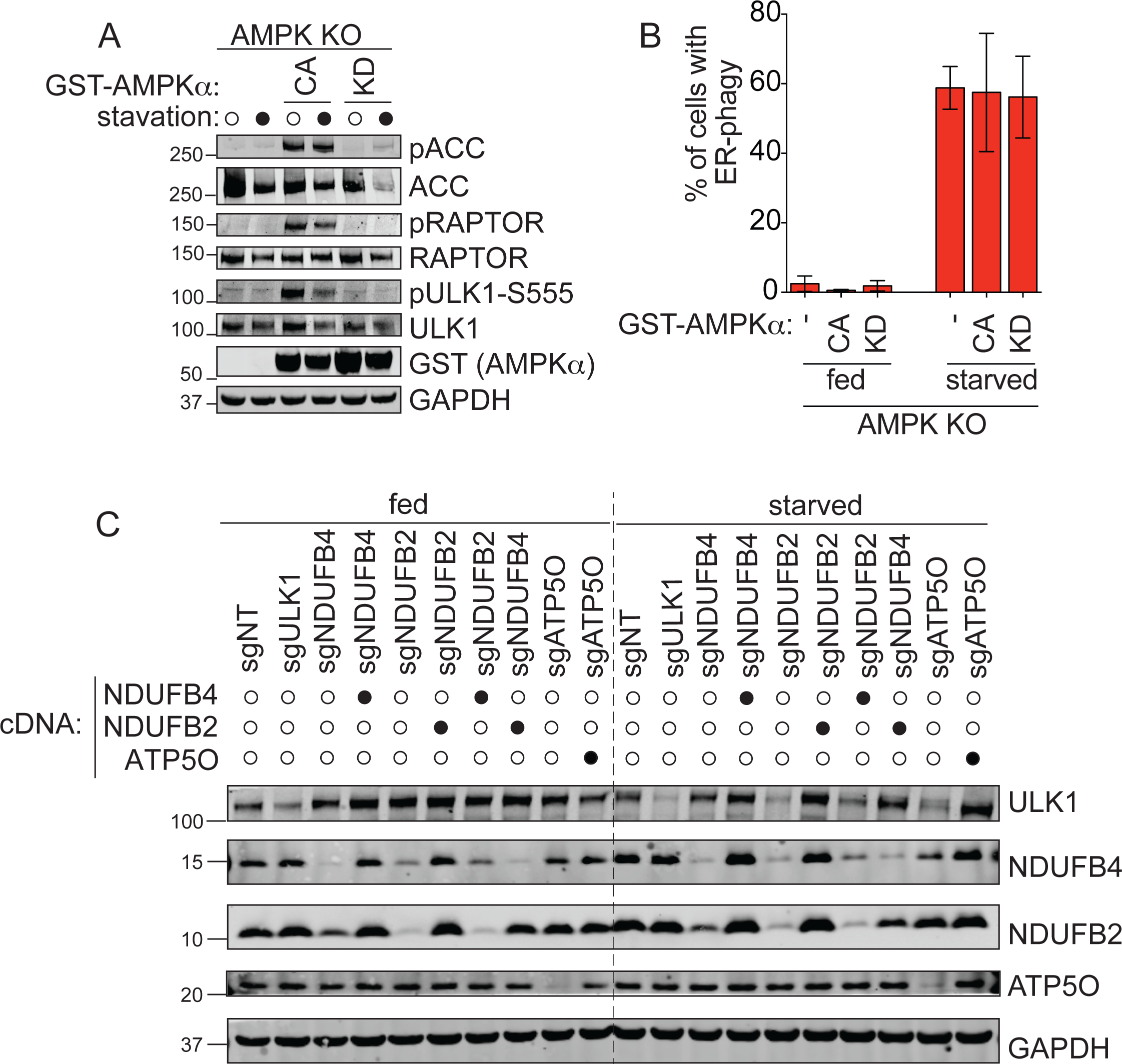
The OXPHOS pathway regulates ER-phagy via ULK1. A. AMPK CRISPR-Cas9 knockout and re-expression. Phosphorylation status of downstream AMPKα targets were analyzed to verify the catalytically active (CA) and kinase dead (KD) AMPKα constructs. AMPKα knockout HCT116 EATR clone from Figure S3G were starved for 16 hours and samples were lysed for Western blotting and immunoprobed for the indicated proteins.
B. Catalytically active (CA) AMPKα does not inhibit ER-phagy. AMPKα knockout HCT116 EATR cells stably expressing catalytically active (CA) AMPKα, or kinase dead (KD) AMPKα were starved for 16hr before FACS measurement of ER-phagy. Data presented as mean ± SD of three biological replicates.
C. ULK1 protein levels are reduced during starvation in NDUFB4, NDUFB2, and ATP5O knockdown cell lines. HCT116 CRISPRi EATR cells were transduced with NDUFB4, NDUFB2, or ATP5O cDNA constructs, and then transduced with sgRNAs targeting ULK1, NDUFB4, NDUFB2, or ATP5O. Cells were lysed for Western blotting and immunoprobed for the indicated proteins.

Unexpectedly, we instead found that interfering with OXPHOS reduces levels of the autophagic kinase ULK1 under conditions that promote ER-phagy. Knockdown of NDUFB2, NDUFB4, and ATP5O did not affect ULK1 under basal conditions, but during starvation the knockdown of OXPHOS components reduced levels of ULK1 almost as much as knockdown of ULK1 itself (Fig 4C), with ULK1 transcript levels moderately affected (Fig S3H). Cognate cDNA re-expression in the appropriate stable knockdown background rescued levels of ULK1, and these same conditions also rescued ER-phagy (Fig 4C, 2D-E). Expressing a non-cognate OXPHOS cDNA in a mismatched knockdown background did not rescue levels of ULK1 nor the ability to perform ER-phagy (Fig 4C, 2D). General autophagy proceeds normally when OXPHOS components are knocked down (Figure 2C, S2C), and so the amount of ULK1 remaining is apparently sufficient to support general autophagy but not ER-phagy. These data reveal an unanticipated mode of regulation for ULK1 and imply a prioritization of general autophagy over ER-phagy during severe energy stress (Fig. S3I).

### DDRGK1-mediated UFMylation regulates ER-specific autophagy

The genome-wide screen for ER-phagy regulators yielded several hits that are localized to the ER and/or involved in ER-related processes ER (Fig 1E, 5A). We focused on one of these factors, DDRGK1/C20orf116/UFBP1, which has emerging roles in ER homeostasis (Leto et al., 2019; Liu et al., 2017; Walczak et al., 2019).

Individual, stable knockdown of DDRGK1 resulted in inhibition of starvation-induced ER-phagy (Fig 5B-C, S4A), but had no effect on general autophagy (Fig 5C-D). Immunofluorescence confirmed that an mCherry-tagged DDRGK1 construct co-localized with the ER (Fig. 5E). DDRGK1 is reported to be post-translationally modified by UFMylation, and which is in turn required for further UFMylation of other factors (Cai et al., 2015; Liu et al., 2017; Wei and Xu, 2016; Wu et al., 2010). UFMylation involves the sequential activation, conjugation and ligation of UFM1 to a target substrate via an E1 (UBA5), E2 (UFC1), and E3 (UFL1) cascade that mirrors ubiquitin conjugation (Fig S4B) (Daniel and Liebau, 2014; Komatsu et al., 2004). We found that stable knockdown of UFL1 led to decreased DDRGK1 protein levels in a proteasome-dependent manner and also inhibited ER-phagy (Fig. 6A-B, Fig S4C), as did knockdown of the UFM1 modifier (Fig. 6C). Double knockdown of both UFL1 and DDRGK1 did not result in greater inhibition of ER-phagy as compared to individual depletion of either factor, suggesting that UFL1 and DDRGK1 act in the same pathway to regulate ER-phagy (Fig S4D – E). Stable re-expression of UFL1 in UFL1-depleted cells rescued levels of DDRGK1 and restored ER-phagy (Fig 6D-E), but over-expression of DDRGK1 in UFL1-depleted cells led to high levels of DDRGK1 without ER-phagy (Fig 6D-E). Hence, DDRGK1-dependent UFMylation is a key mediator of ER-phagy.

**Figure 5.**
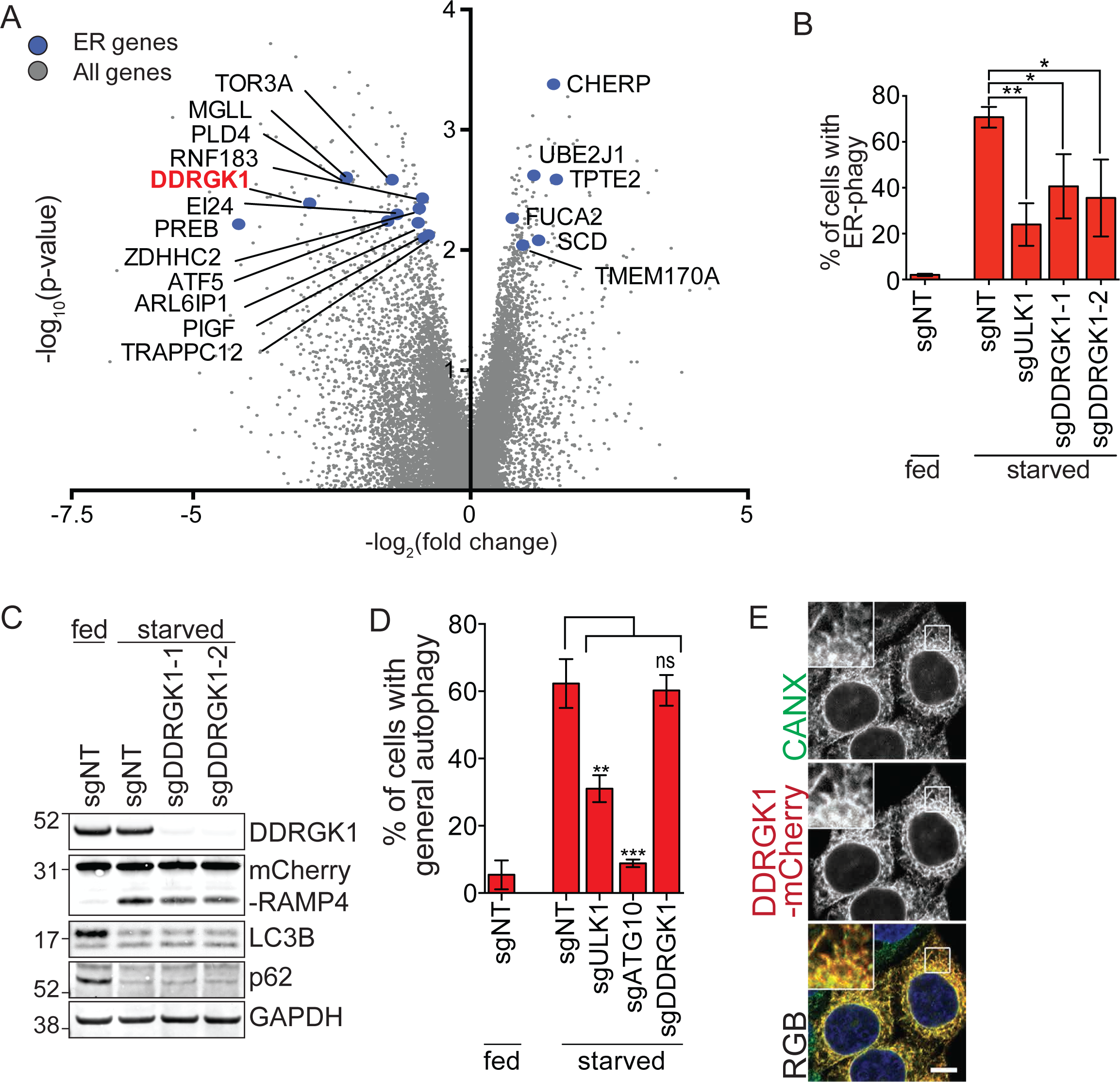
DDRGK1 is specifically required for ER-phagy. A. ER-phagy CRISPRi screen identifies genes that are functionally or physically associated with the ER. DDRGK1 is highlighted in bold and red.
B. DDRGK1 depletion results in inhibition of ER-phagy based on EATR assay. HCT116 CRISPRi EATR cells were transduced with sgRNAs targeting ULK1 or DDRGK1 and starved for 16hr before FACS measurement for ER-phagy. Data presented as mean ± SD of three biological replicates. P value indicates two-tailed unpaired t-test (*, P < 0.05, **, P < 0.01).
C. CCER assay shows ER-phagy inhibition upon DDRGK1 knockdown. HCT116 CRISPRi CCER cells were transduced with sgRNAs targeting DDRGK1 and starved for 16hr. Cells were lysed for Western blotting of the indicated proteins.
D. DDRGK1 knockdown does not affect general autophagy. HCT116 CRISPRi cells stably expressing eGFP-mCherry-LC3B constructs were transduced with sgRNAs targeting either ULK1, ATG10 or DDRGK1. Cells were starved for 16hr before FACS measurement for general autophagy. Data represents mean ± SD of three biological replicates. P value indicates two-tailed unpaired t-test (**, P < 0.01, ***, P<0.001).
E. DDRGK1 localizes to the ER. HeLa cells were stably transduced with DDRGK1-mCherry construct and immunostained for calnexin (CANX) as an ER marker. Insets represent three-fold enlargement of boxed areas. Scale bar represents 10µm.

**Figure 6.**
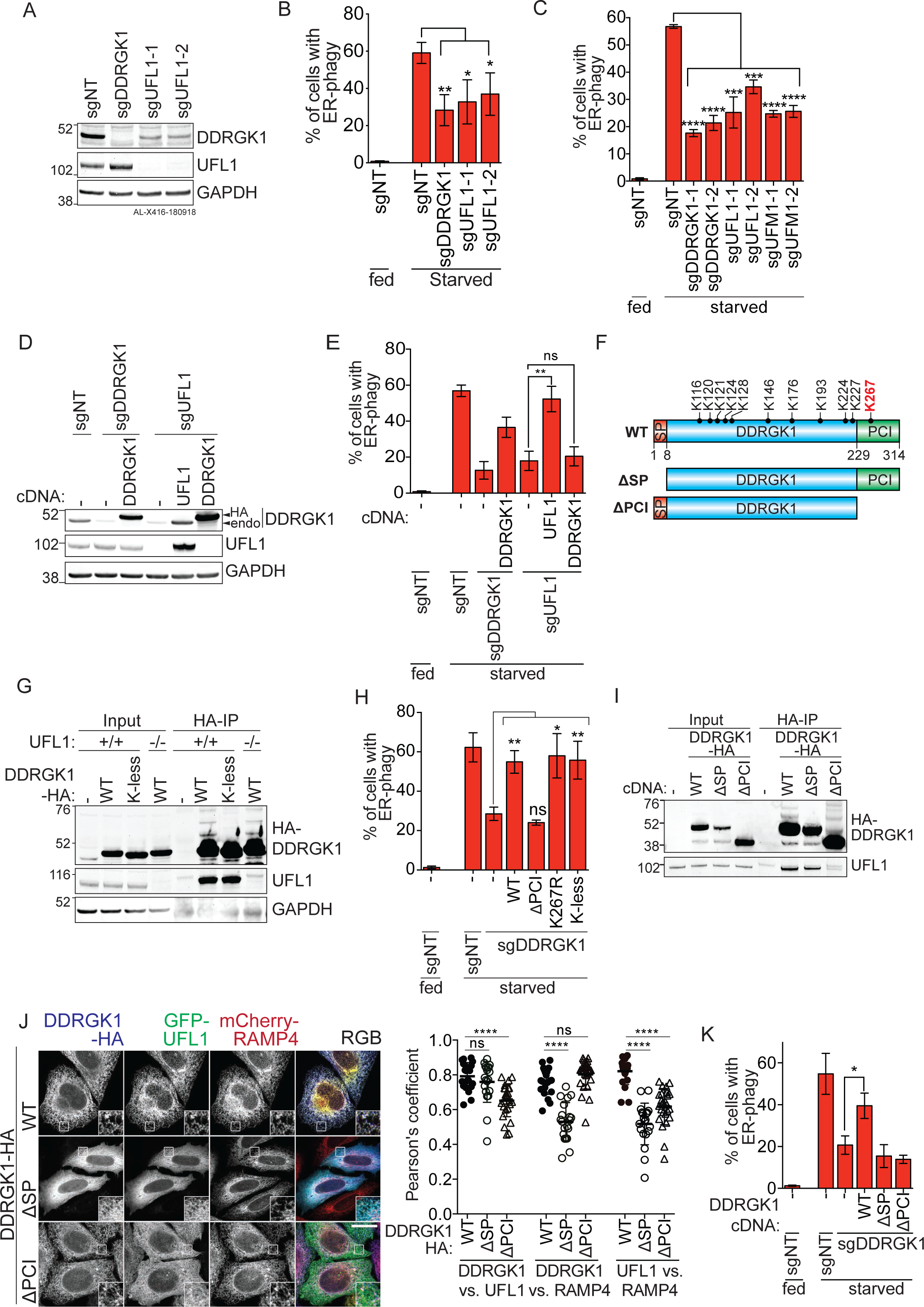
ER-phagy is mediated by DDRGK1-dependent UFMylation and ER localization. A. UFL1 knockdown results in decreased DDRGK1 protein levels. HCT116 CRISPRi EATR cells were transduced with sgRNAs targeting either DDRGK1 or UFL1. Cells were harvested for Western blotting to assess the effect of UFL1 knockdown of DDRGK1 protein levels.
B. UFL1 knockdown phenocopies DDRGK1 knockdown and results in ER-phagy inhibition. The cells generated in (A) were starved for 16hr before FACS measurement for ER-phagy. P value indicates two-tailed unpaired t-test (*, P < 0.05, **, P<0.01).
C. UFMylation components, including UFL1, DDRGK1 and UFM1 are required for ER-phagy. HCT116 CRISPRi EATR cells expressing the indicated sgRNAs were starved for 16hr before FACS measurement for ER-phagy. P value indicates two-tailed unpaired t-test (***, P < 0.001, ****, P<0.0001).
D. Re-expression of UFL1 restores DDRGK1 protein levels. HCT116 CRISPRi EATR cells were transduced with sgRNAs targeting either DDRGK1 or UFL1. Cells were then further transduced with either DDRGK1-HA or HA-UFL1. Cells were harvested for Western blot analysis of DDRGK1 and UFL1 protein levels.
E. Re-expression of DDRGK1 in UFL1 knockdown cells does not rescue ER-phagy. The cell lines generated in (E) were starved for 16hr and then subjected to FACS measurement for ER-phagy. Data represents mean ± SD of three biological replicates. P value indicates two-tailed unpaired t-test (**, P<0.01).
F. Schematic of DDRGK1 domains and conserved Lysine residues. There are twelve conserved Lysine residues on DDRGK1. The reported Lysine residue that is a major site for UFMylation (K267) is labelled in red. Also shown are the two truncated forms of DDRGK1 that either lacks the N-terminal signaling peptide (ΔSP) or the C-terminal proteasome component (ΔPCI).
G. Post-translational modification of DDRGK1 occurs on lysine residues. Parental or UFL1 knock-out HCT116 cells were transfected with either wildtype (WT) DDRGK1-HA or the Lysine-less (K-less) DDRGK1-HA constructs. Cells were then harvested for immunoprecipitation of the HA-epitope and the higher molecular weight species of DDRGK1 is resolved by Western blotting.
H. DDRGK1’s role during ER-phagy does not require post-translational modification on any lysine residue. HCT116 CRISPRi EATR cells were transduced with DDRGK1 sgRNA and then rescued using the indicated DDRGK1-HA mutant constructs. Cells were then starved for 16hr and ER-phagy was measured by FACS analysis. Data represents mean ± SD of three biological replicates. P value indicates two-tailed unpaired t-test (*, P< 0.05, **P<0.01).
I. DDRGK1 interacts with UFL1 via DDRGK1’s PCI domain. Parental HCT116 cells were stably transfected with the indicated DDRGK1-HA mutant constructs. Cells were then harvested for HA-immunoprecipitation to determine the DDRGK1 domains that are required for UFL1 interaction.
J. DDRGK1 recruits UFL1 to the ER. DDRGK1 knock-out HeLa cells were stably transduced with mCherry-RAMP4 (ER marker) and the indicated DDRGK1-HA mutant constructs. Cells were then transiently transfected with GFP-UFL1 for 24hr. Cells were then fixed and immunostained for HA epitope. Representative images are shown. Insets represent three-fold enlargement of boxed areas. Scale bar represents 10µm. Pearson’s Correlation coefficient was measured between DDRGK1 vs. UFL1, DDRGK1 vs. RAMP4, and UFL1 vs. RAMP4. Data was generated from one biological replicate and 20-26 cells were analysed from each condition. P-value indicates two-tailed unpaired t-test (****, P<0.0001).
K. DDRGK1’s role during ER-phagy requires both the SP and PCI domains. HCT116 CRISPRi EATR cells with DDRGK1 knockdown were rescued using the indicated DDRGK1-HA mutant constructs. Cells were then starved for 16hr and ER-phagy was measured by FACS analysis. Data represents mean ± SD of three biological replicates. P value indicates two-tailed unpaired t-test (*, P< 0.05).

DDRGK1 is reported to be UFMylated by UFL1 on one or more lysines and thereby stabilized (Fig 6F) (Wu et al., 2010). Using immunoprecipitation of DDRGK1 point mutants, we indeed found higher molecular weight species consistent with lysine post-translational modification of DDRGK1 (Fig 6G, S4F). However, this modification was unaffected by CRISPR-Cas9 knockout of UFL1 or knockdown of UFM1 (Fig 6G and S4F). Furthermore, knockdown of UFM1 had no effect on the abundance of DDRGK1 (Fig S4G), and DDRGK1 still stably interacted with UFL1 even when all twelve conserved lysines were mutated (Figure 6G). Taken together, these data indicate that the stability of endogenous DDRGK1 is maintained not by UFMylation, but by its interaction with UFL1. Along these lines, we found that DDRGK1’s ability to promote ER-phagy was independent of its major reported site of UFMylation on Lys267 (Fig 6H & S4H). Eleven other lysine mutants could also substantially support ER-phagy (Fig S4I-J), as could a DDRGK1 mutant with all twelve conserved lysines mutated (Fig 6H & S4H).

Using immunoprecipitation and immunofluorescence, we found that UFL1 interacts strongly with DDRGK1 and is localized to the ER (Fig 6I & J, S5C). Deleting DDRGK1’s N-terminal ER-targeting signal peptide (SP) still supported the DDRGK1-UFL1 interaction, but led to cytoplasmic localization of both DDRGK1 and UFL1 and did not support ER-phagy (Fig 6I-K, S5A-C). Removing DDRGK1’s C-terminal PCI protein-interaction domain led to normal ER localization of DDRGK1 (Fig 6J, S5B), but abolished its interaction with UFL1 (Fig 6I, S5C). This led to cytoplasmic localization of UFL1 and abolished the cell’s ability to perform ER-phagy (Fig 6J & K, S5C).

Overall, we found that DDRGK1-dependent ER-phagy is not mediated by UFMylation of DDRGK1, as has been reported for DDRGK1’s involvement in UPR and other signaling pathways (Lemaire et al., 2011; Yoo et al., 2014). Instead, ER-phagy is mediated by a functional interaction between DDRGK1 and UFL1 that recruits UFL1 to the ER. This suggested that UFMylation of some downstream ER-resident target(s) mediates ER-phagy. UFMylation is reported to stabilize the proteins it modifies (Cai et al., 2015; Egunsola et al., 2017; Liu et al., 2017; Wu et al., 2010; Yoo et al., 2014), and we knocked down previously reported substrates of UFL1, but found that none of them affected ER-phagy (Fig S5D-E).

We likewise found no evidence for DDRGK1-mediated UFMylation and stabilization of IRE1α (Fig S6A), which was previously reported to promote the UPR (Liu et al., 2017). However, we conversely found that depletion of DDRGK1, UFL1, or UFM1 in multiple cell types resulted in elevated levels of IRE1α (Fig 7A, S6B-D). These results suggested that an inability to perform UFMylation-dependent ER-phagy could lead to upregulation of an ER stress response through IRE1α, which senses misfolded proteins in the ER lumen. Indeed, knockdown of UFMylation ER-phagy factors led to elevated levels of several other UPR proteins including PERK, BiP and CANX (Fig 7A, S6B). We also observed an increase in levels of CLIMP63 (ER sheet marker) and REEP5 (ER tubule marker), suggesting ER expansion (Fig 7A, S6B) (Schuck et al., 2009). Consistent with this idea, immunofluorescence of DDRGK1 CRISPR-Cas9 knockout HeLa cells showed increased CANX staining (Fig S5D-E). Knockdown of DDRGK1, UFL1, or both DDRGK1 and UFL1 led to modest but consistent transcriptional upregulation of the UPR transcripts PERK and BiP, increased differential splicing of XBP1, and transcriptional upregulation of CLIMP63 and REEP5 (Fig 7B). IRE1α showed higher protein levels upon DDRGK1 knockdown but no change in transcript abundance (Fig 7A-B), indicating that IRE1α protein levels could be post-translationally regulated in response to UFM1 signaling. The transcriptional upregulation of multiple UPR transcripts, differential splicing of XBP1, post-translational upregulation of IRE1α, and ER expansion are all consistent with increased ER stress and consequent UPR under conditions where UFMylation-dependent ER-phagy cannot be executed.

**Figure 7.**
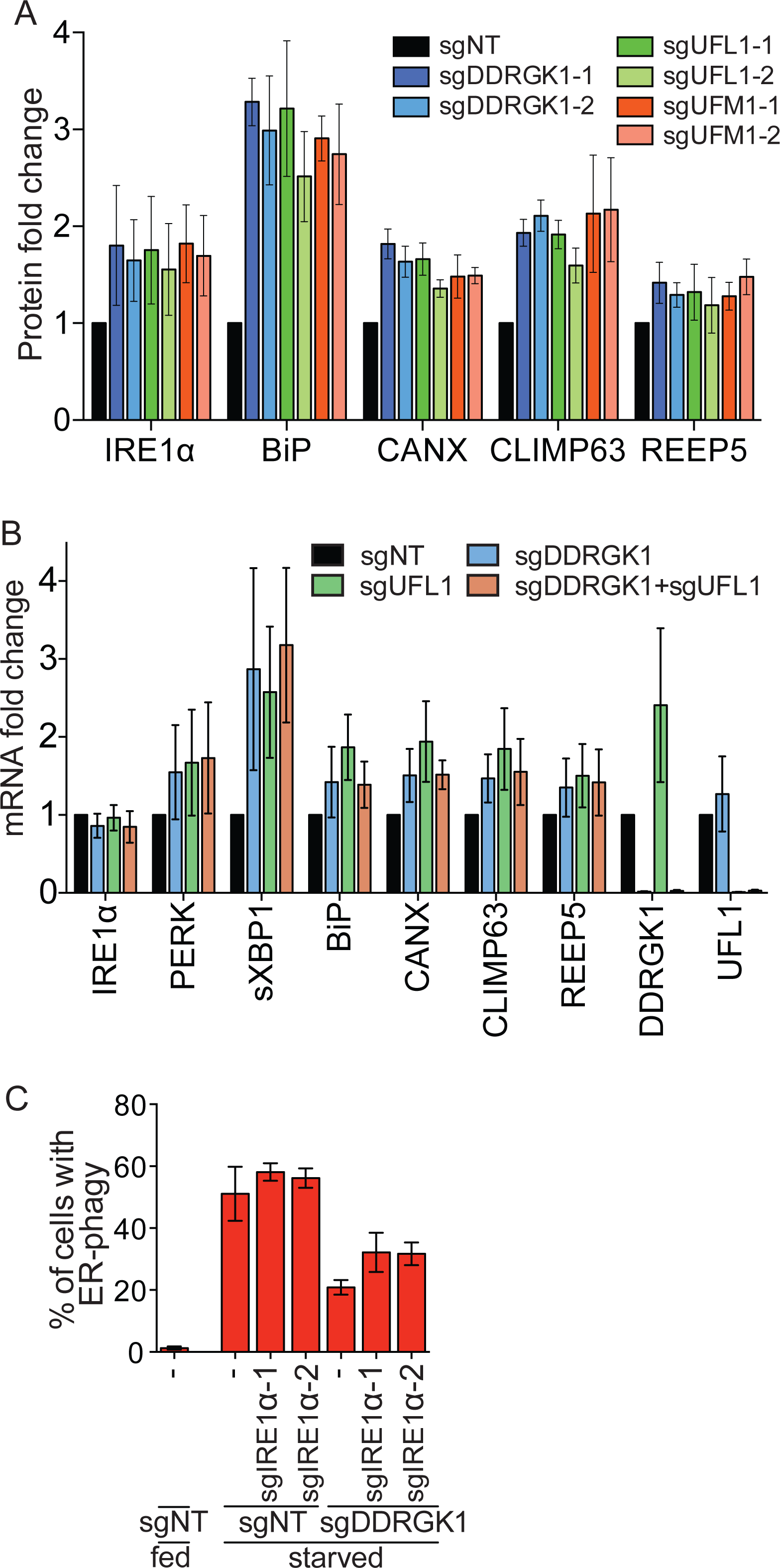
UFMylation-mediated ER-phagy represses IRE1α UPR. A. Knockdown of DDRGK1, UFL1 or UFM1 results in upregulation of UPR and general ER protein levels. HCT116 cells were transduced with sgRNAs targeting DDRGK1, UFL1 and UFM1 (two sgRNAs per gene). Cells were harvested for Western blot analysis of the depicted proteins. Graph represents densitometry measurement of the indicated proteins upon protein knockdown. Data represents mean ± SD of three biological replicates. A representative blot is shown in Fig S6B. (Note that Fig S6B is an expansion of Fig S4G).
B. Knockdown of UFL1 and/or DDRGK1 causes transcriptional upregulation of UPR and ER markers, except IRE1α. HCT116 CRISPRi cells were transduced with sgRNA targeting DDRGK1, UFL1 or both. Cells were then harvested for qRT-PCR measurement of the indicated ER or UPR genes. Data represents mean ± SD of three biological replicates.
C. Knockdown of IRE1α partially restores ER-phagy in DDRGK1 depleted cells. HCT116 CRISPRi EATR cells transduced with the indicated sgRNAs and then starved for 16hr before FACS measurement for ER-phagy. Data represents mean ± SD of three biological replicates.

IRE1α senses unfolded proteins in the ER lumen and so is a good candidate to mediate stress signals caused by defective ER-phagy. We did not observe direct UFMylation of IRE1α under either fed or starved conditions (Fig S6A). But knockdown of IRE1α in DDRGK1 depleted cells reversed the high levels of ER stress markers caused by an inability to execute ER-phagy (Fig S6F). Knockdown of IRE1α had only a modest reciprocal effect upon ER-phagy, and only somewhat reversed the ER-phagy defect induced by loss of DDRGK1 (Fig 7C). Hence, UFMylation-induced ER-phagy is upstream of IREα signaling, further indicating that the loss of ER-phagy induces ER stress through the accumulation of unfolded proteins that are sensed by IRE1α.

Overall, our data indicate that DDRGK1-mediated, ER-resident UFMylation through UFL1 is critical for ER-phagy. DDRGK1 recruits UFL1 to the ER surface, analogous to the role of PINK1 substrates in recruiting Parkin to the mitochondria during mitophagy (Genschik et al., 2013; Léon and Haguenauer-Tsapis, 2009). We propose that the inability to UFMylate downstream ER substrates leads to an inability to execute ER-phagy, resulting in the consequent build-up of ER stress, and eventual activation of the unfolded protein response via IRE1α (Fig S6G).

## Discussion

ER-phagy is a new and relatively unexplored branch of ER quality control that sends entire portions of the ER for destruction in the lysosome. Beyond a handful of ER-phagy receptors and ER remodeling proteins, knowledge of the logic and factors that regulate ER-phagy is still in its infancy. Our genome-wide ER-phagy screen provides a rich set of genes and pathways that are involved in ER-phagy. Among these, we identified several aspects of the core autophagy machinery, consistent with studies showing that ER-phagy shares effectors with general autophagy (Grumati et al., 2017; Khaminets et al., 2015; Smith et al., 2018). In particular, autophagosomal membrane activation and expansion separates ER-phagy from ERLAD, as the latter overrides the need for autophagosomal engulfment and instead forms ER-derived vesicles that directly fuse with lysosomes (Fregno et al., 2018). Moreover, ER-phagy and ERLAD pathways are activated by different upstream signals of starvation and proteasome-resistant protein aggregation, respectively.

So far, nutrient starvation is the only stress known to induce ER-phagy. We found that ER stress-inducing compounds do not lead to ER-phagy, but repression of ER-phagy by knockdown of DDRGK1 UFMylation induces ER stress and the UPR. UFMylation has previously been linked to ER stress through unclear mechanisms, which we now connect to inhibition of ER-phagy (DeJesus et al., 2016; Leto et al., 2019; Walczak et al., 2019). How is it that ER stress does not induce ER-phagy, but an inability to perform ER-phagy induces ER stress? Under nutrient depletion, protein misfolding may increase in the ER, but these signals are repressed as cells catabolize the protein- and lipid-rich organelle. Blocking ER-phagy could then result in the toxic accumulation of excessive ER and misfolded ER-resident proteins that cannot be sufficiently kept in check by ERAD, thus activating the UPR. Under this model, blocking ER-phagy leads to UPR as a byproduct of ER stress that is no longer relieved by eating portions of the ER. This hypothesis and ordered prioritization of ER stress-relief pathways will require a great deal of investigation but could lead to a molecular rationale for why cells go to the extreme of ER-phagy.

We found extensive interplay between the mitochondria and ER-phagy. The ER and mitochondria are known to crosstalk at membrane contact sites, including transfer and expansion of the lipid bilayer, Ca^2+^ homeostasis, and mitochondria division (Friedman et al., 2011; Lombardi and Elrod, 2017). The interactions previously described between the two organelles all indicate a regulatory role of ER processes towards mitochondrial homeostasis. We found that impairment of mitochondrial OXPHOS represses ER-phagy, demonstrating that mitochondrial metabolism can also inform decisions in the ER. It remains to be seen whether this ULK1-mediated communication is directly orchestrated via mitochondria-ER contacts or indirectly as a result of metabolic products. Alternatively, inhibition of OXPHOS could initiate UPR that takes over to repress last-resort ER-phagy. Consistent with this idea, a recent screen for regulators of IRE1α found that knockdown of mitochondrial metabolism genes stimulates the UPR, though the mechanism remains to be determined (Adamson et al., 2016). Cellular energy levels are regulated by multiple energy sensing mechanisms that have complex roles during general autophagy (Egan et al., 2011; Herzig and Shaw, 2018; Kim et al., 2011). The interplay between mitochondrial metabolism and ER homeostasis will no doubt involve a similarly rich set of pathways.

We found that DDRGK1-mediated UFMylation at the ER surface is a key regulator of ER-phagy. Post-translational modifications of an organelle’s surface are widely involved in organelle autophagy. For example, ubiquitylation of PEX5 serves as a signal for peroxisomal autophagy (Nordgren et al., 2015; Zhang et al., 2015), and ubiquitination of multiple mitochondrial substrates promotes mitophagy (Chan et al., 2011; Karbowski and Youle, 2011). Since DDRGK1 recruits UFL1 to the ER surface and their combined ER-resident activity with UFM1 are all required for ER-phagy, we speculate that UFMylation of ER surface protein(s) serves as an effector of ER-phagy. This would be strikingly similar to PINK1’s recruitment of the ubiquitin ligase Parkin to modify multiple mitochondrial surface proteins and initiate mitophagy (Chan et al., 2011; Glauser et al., 2011; Karbowski and Youle, 2011; Wang et al., 2011). UFMylation may play multiple cellular roles, and we have ruled out several previously reported UFMylation targets in the regulation of ER-phagy, though more substrates are still being uncovered (Walczak et al., 2019). In the case of mitophagy, a single causative ubiquitination substrate has remained elusive, and it remains to be seen if this will be the case for UFMylation-dependent ER-phagy.

While defects in ER-phagy have not been explicitly linked to human disease, we note that human mutations in ER-phagy genes such as FAM134B and Atlastins are associated with hereditary neuropathies in OMIM and ClinVar (Abel et al., 2004; Amberger et al., 2015; Kurth et al., 2009). This is similar to the mouse phenotype of FAM134B knockout (Khaminets et al., 2015). Several of the ER-phagy genes we identified by genome-wide screening are also associated with human neurodegenerative phenotypes with previously unclear mechanistic bases, such as Leigh Syndrome (mitochondrial OXPHOS, including ETC chaperones) (Lake et al., 2016), spastic paraplegia (ARL6IP1) (Novarino et al., 2014), encephalopathy (TRAPPC12) (Milev et al., 2017), spinocerebellar ataxia and encephalopathy (UBA5) (Daida et al., 2018; Mignon-Ravix et al., 2018), and severe early-onset encephalopathy and progressive microcephaly (UFC1, UFM1) (Nahorski et al., 2018). It is premature to broadly link deficits in ER-phagy to human disease, but the similar phenotypes stemming from mutations in different ER-phagy factors is provocative. Our work lays the foundation for future understanding of ER-phagy and its interplay with the ER stress response, as well as the consequences of ineffective ER-phagy. Further mechanistic dissection of the 200 high-confidence ER-phagy regulators and executors identified here will hopefully shed light on this dramatic process.

## Methods

### Design, Production and titering of sgRNA library lentivirus

The genome-wide CRISPRi-V2 library was a gift from the Weismann lab (Addgene catalog #1000000093) and contains 5 sgRNAs per gene. For the pilot autophagy screen, we designed a comprehensive sgRNA library that targets all the reported TSS (10 gRNAs per TSS) of 31 genes that are involved in general autophagy. Overall, a total of 3301 gRNAs were designed (Table S1). The protospacer oligos were annealed and ligated to pCRISPRia vector (Addgene 84832) according to the protocol established by the Weissman lab (https://weissmanlab.ucsf.edu/CRISPR/Pooled_CRISPR_Library_Cloning.pdf) (Horlbeck et al., 2016). In addition, we added in 10% of a custom built non-targeting sgRNA library prior to virus production.

The following paragraph describes the transfection protocol for one 15 cm plate of HEK293T cells. On Day 0, 7.5 × 10^*6* HEK293T cells were seeded in a 15 cm plate in 20 mL of DMEM medium with 10% FBS. The following day HEK293T cells were transfected. In a 15 mL tube, 2.8 mL of Opti-MEM was mixed with 90 µL of Mirus LT1 transfection reagent and incubated at room temperature for 5 minutes. In an eppendorf tube, 12 µg of delta VPR, 3 µg of VSVG, and 15 µg of library plasmid were combined. The plasmids were then added to the Opti-MEM and Mirus mixture and incubated at room temperature for 20 minutes. The media was changed the following day. On Day 3, the virus was harvested using a 0.45 µm syringe filter, aliquoted into 1 mL tubes, and snap frozen. If more than one 15 cm plate of virus was produced for one library, the virus across those plates were pooled and mixed prior to aliquoting into eppendorf tubes. Virus was harvest on Day 4 as well.

Next, the virus was titered to determine the infectivity of the virus in the HCT116 cells. HCT116 were plated in a series of 6 well plates such that each well had cells and there was one 6 well plate per sub-library per time point (i.e. 48 or 72 hour virus harvest). One well on each plate was not transduced with any virus. The virus was titered such that is diluted 2-fold, 4-fold, 8-fold, 16-fold, and 32-fold. Polybrene was used at a concentration of 8 µg/mL. Fresh media were replaced 24hr post transduction. The cells were harvested 48 hours post viral transduction for flow cytometry and the percentage of BFP positive cells was recorded. The optimal virus dilution is defined as dilution-fold that results in less than 20% of BFP positive cells.

### CRISPRi screen: cell generation, virus transduction, puro selection, and sort

HCT116 cells expressing a dcas9-KRAB and EATR reporter was constructed as described previously (Liang et al., 2018). The library contained seven unique sub-libraries and each sub-library was transduced separately, such that each sgRNA had an average of 500x coverage after transduction (Day 1). Puromycin selection for positively-transduced cells was performed 48 hours post transduction (Day 3). On Day 7, the sub-libraries were pooled proportionally based on the number of sgRNAs and cells were maintained at 500x coverage. On Day 10, cells were treated with doxycycline (4µg/ml) for 16 hours to induce EATR expression and on Day 11, cells were treated with EBSS for 16 hours. Cells were then collected for sorting – cells were gated into the 25% of cells with most ER-phagy and 25% of cells with the least ER-phagy. A background population of cells was collected for downstream NGS analysis of relative enrichment.The entire CRISPRi screen was performed in two biological replicates.

### NGS Sample Preparation and screen analysis

Genomic DNA was harvested using the Macherey-Nagel gDNA extraction protocol. The background samples required the XL kit whereas the midi kit was sufficient for sorted cells. After elution, the genomic DNA was treated with SbfI-HI restriction enzyme and incubated overnight at 37 □ to liberate the DNA fragment encoding the sgRNA sequences.

Samples were run an agarose gel and the gel piece around the 500 bp size (region containing the sgRNA sequence) was excised. The gel was melted in 55 □ water bath and 1/100 by volume of 3 M NaAc (pH 5.2) was added to each tube and then solution was passed through an MN column. Each column was washed twice with NT3 buffer. The column was incubated for 5 minutes in 20 µL of heated elution buffer (98□) and then spun. The elution step was repeated so that the final elution volume was 40 µL.

A standard PCR protocol was used with Phusion High Fidelity Enzyme and 3% DMSO final concentration.The forward primer contained a TruSeq Index that would be subsequently used during NGS analysis. Before proceeding with a full scale PCR of the samples, a test PCR for each sample was run to determine the proper number of cycles (21, 23, or 25 cycles). The cycle number was identified individually for each sample that allowed a visible band on a TBE gel after staining with ethidium bromide, but not an oversaturated PCR product that could compromise the representation of gRNAs within the sample.

After the optimal cycle number was determine, a total of twelve 100 µL PCRs were done with 3 µL of template per reaction (from the abovementioned elution). The forward primer contained a TruSeq Index that would be subsequently used during NGS analysis. After completion of the PCR, the twelve reactions were pooled together and mixed. 300 µL of the pooled PCR was taken for subsequent PCR clean-up.

195 µL of SPRI beads was added to the pooled PCR and incubated at room temperature for 10 minutes. The samples were attached to a DynaMag for 5 minutes. The supernatant (which has the sample) was transferred to a new tube. 300 µL of SPRI beads were added and incubated for another 10 minutes. The samples were attached to a DynaMag for 5 minutes and the supernatant was discarded (samples attached to the beads). The beads were washed twice with 80% ethanol. After removal of the last supernatant, the beads were spun down, and excess ethanol was removed. The samples were air dried for 10 minutes and resuspended in 35 µL of water. DNA concentration was quantified using Qubit Fluorometric Quantitation (Thermo Fisher Scientific) and the samples were pooled proportionally to cell number and sequenced on a HiSeq 2500 such that each sgRNA sequence was covered at least 30 times.

Screening data was analyzed using standard protocols in MaGECK and ScreenProcessing (Horlbeck et al., 2016; Li et al., 2014, 2015). MaGECK was used for the pilot autophagy library, while ScreenProcessing was used for the genome-wide library. Briefly, gRNAs were quantified in each pool of cells based by matching reads back to the appropriate library reference, each pool was normalized by total number of reads, and gRNA distributions were compared to the background. Non-targeting gRNAs were explicitly used in each software package. MaGECK and ScreenProcessing integrate multiple gRNAs into gene-level phenotypes (e.g. log_2_-fold-change) and p-values using different approaches (Horlbeck et al., 2016; Li et al., 2014, 2015).

### sgRNA plasmid cloning procedures for individual plasmids

The sgRNA sequences for genome-wide screening were based on the Weissman CRISPRi-v2 library and contained 5 sgRNAs per gene. The sgRNA sequences for autophagy-related genes used for the pilot test run were custom-designed to target all reported transcription start site (TSS) of each gene and contained 10 sgRNAs per TSS. sgRNA plasmids were cloned by annealing and ligating sgRNA-containing short oligos to the CRISPRi-v2 vector (addgene 84832) via the previously described protocol (Horlbeck et al., 2016). Knockdown efficiency of each guide was measured either by western blot or qRT-PCR. All sgRNA constructs used in this study are detailed in Table S3.

### shRNA plasmid cloning for DDRGK1

Short hairpin RNA (shRNA) was used to knockdown DDRGK1 in cell lines that do not express dCas9-KRAB constructs. Briefly, non-targeting (5’-CCTAAGGTTAAGTCGCCCTCG-3’) and DDRGK1-targeting (5’-GGCTCTGCTAGTCGGCTTTAT-3’) shRNAs were cloned into pLKO.1 puro construct (Addgene #8453) according to protocol described in Addgene (https://www.addgene.org/tools/protocols/plko/?gclid=Cj0KCQiAm5viBRD4ARIsADGUT25ZCGNPeQSFvLqSwvg2tHDkCc9zOZsLdaUffZzNTRYzI_YOlKFVQdUaAqbfEALw_wcB)

### cDNA plasmid cloning procedures

The open reading frame (ORF) of the constitutively active-AMPK construct (CA-AMPK) was sub-cloned from addgene #27632 (Egan et al., 2011). The AMPK kinase dead (AMPK-KD) plasmid was a Lys-to-Arg mutation at position K47R (AAG to CGG) generated by extension PCR followed by Gibson Assembly (Gibson et al., 2009). The pBMN-YFP-Parkin construct was from Addgene (Addgene #59416). Unless stated otherwise, all remaining ORFs described in this article were obtained from PCR amplification of pooled HCT116 cDNA. The ORFs were cloned into pLenti-XI destination vector with neomycin resistance. Briefly, an original pLenti-X1-Neo-eGFP-LC3B vector was first digested with restriction enzymes BamHI and XbaI to remove the insert. Then, Gibson Assembly was used to insert the gene-of-interest and the desired epitope or fluorescent tag into the pLenti-X1 vector. All overexpression constructs used in this study are detailed in Table S4.

### Cell culture

Cells were cultured at 37 **°**C with 5% CO_2_ in a humidified atmosphere. All cells were cultured in DMEM-GlutaMAX medium supplemented with 10% FBS, 0.1 mM non-essential amino acids (Gibco), 1 mM sodium pyruvate (Gibco), 100 U/mL penicillin (Gibco), and 100 g/mL streptomycin (Gibco). Cell lines were obtained from the Berkeley Cell Culture Facility and were verified mycoplasma free with MycoAlert Mycoplasma Detection Kit (Lonza).

### Lentiviral packaging and transduction

Lentiviral packaging was performed in HEK293T cells using either TransIT-LT1 Transfection Reagent (Mirus) or Lipofectamine 3000 (ThermoFisher Scientific) according to the manufacturer’s protocol. For more details, refer to previously described methods (Liang et al., 2018).

### Knockout Cell Line Generation

AMPK knockout cell lines were generated using Cas9 RNPs and nucleofection as detailed previously (Lingeman et al., 2017). The sgRNA protospacer sequences were validated and used previously by the Shaw lab (Toyama et al., 2016). The protospacer sequences are as follow: AMPKα1- sgRNA1- GGCTGTCGCCATCTTTCTCC; AMPKα1-sgRNA2- GAAGATCGGCCACTACATTC; AMPKα2-sgRNA1- TCAGCCATCTTCGGCGCGCG; AMPKα2-sgRNA2-GAAGATCGGACACTACGTGC. After nucleofection, HCT116 cells were serial diluted into 96 well plates such that there was on average of 0.7 cells/well. AMPK KO clones were screened by Western blotting. DDRGK1 and UFL1 knockout cell lines were generated by transient transfection of two pSpCas9(BB)-2A-GFP (PX458) (Addgene #48138) plasmids carrying sgRNAs that each target the downstream and upstream regions of the transcription start site. The protospacer sequences are as follow: DDRGK1-sgRNA1- ATGAGATCCCGGCCTCAGGG; DDRGK1-sgRNA2- TAGGAGATGCCGCTGCACCA; UFL1-sgRNA1- CTGACTCGCAGTAGACGCGG; UFL1-sgRNA2-GCCTAATT TGGGCTCCACAA. GFP-positive cells were single-cell sorted 48hrs post transfection and DDRGK1/UFL1 KO clones were screened by Western blotting.

### Flow Cytometry Analysis of EATR cells

Flow cytometry of EATR assay was performed using an Attune NxT Flow Cytometer and subsequent analysis was performed using FlowJo 10.1 (Liang et al., 2018). All EATR experiments were performed using live cells to prevent reversal of eGFP quenching post-fixation. The intensities for both eGFP and mCherry of the EATR cells at fed condition were used as references to define the gate for zero ER-phagy events. Following stimulation, ER-phagy detection is based on the shift of cell population into the ER-phagy gate. On average, 5 to 10,000 cells were analyzed per condition and all statistical analyses were performed using data from at least three biological replicates.

### Cell Treatments

ER-phagy was induced with media starvation using EBSS with calcium, magnesium, and phenol red (Invitrogen e10043). For EATR and CCER assays, cells were plated 48 hours prior to EBSS treatment. EATR expression is induced using 4 µg/ml doxycycline 24 hrs prior to starvation. Unless otherwise stated, starvation treatment was carried out for 16 hours. Cells in fed conditions indicate incubation in complete DMEM described above.

For all experiments except the Seahorse assay, rotenone was used at a final concentration of 3 µM, antimycin A was used at a concentration of 0.5 µM, and oligomycin A was used at a concentration of 3 µM. Cells were treated with these drugs in two phases for a total of 40 hours. First, cells were treated for 24 hours with complete DMEM, then immediately treated again for 16 hours in EBSS media or complete DMEM. Epoxomicin and folimycin treatments were co-administered with EBSS starvation at 100 nM final concentration.

### Quantitative real-time PCR (qRT-PCR)

RNA extraction was performed using Directzol RNA miniprep kit (Zymo Research) according to manufacturer’s instruction. 1µg of RNA per sample were used for reverse transcription using SuperScript™ III First-Strand Synthesis System (Invitrogen) according to manufacturer’s instruction. qRT-PCR reaction was set up using Fast SYBR Green Mastermix (Applied Biosystems) and run in triplicates using StepOne Plus Real-Time PCR system (Applied Biosystems) (Liang et al., 2018). A complete list of all primers used are compiled in Table S5.

### Western blotting

To prepare samples for western blot, cells were lysed in RIPA buffer (Millipore), supplemented with Halt Protease Inhibitor Cocktail and Phosphatase Inhibitor Cocktail (both ThermoFisher). Cells were lysed on ice for 30 minutes and spun at 14000 rpm for 10 minutes to remove insoluble debris. Protein concentrations were quantified by Bradford assay. Lysates were normalized based on protein concentration and NuPage LDS Sample Buffer (4x) was added (Invitrogen). Samples were boiled at 98°C for 5 minutes.

Between 40-50µg of samples were run on NuPAGE Bis-Tris 4–12% gels in NuPage MES SDS Buffer (Invitrogen) for 40 minutes at 200 V and transferred to 0.4-µm nitrocellulose membranes using a semi-dry transfer system (Bio-Rad Catalog #1704150) at 1.3 A and 25 V for 15 minutes. After transfer, membranes were blocked with 5% (w/v) milk in TBS-T for 30 minutes, and subsequently washed with TBS-T three times. Primary antibodies were diluted at the appropriate concentration in 5% BSA (w/v) in TBS-T. The membrane was incubated in primary antibody for either 1-2 hours at room temperature or overnight at 4°C. The membrane was washed with TBST three times for five minutes each. The blots were incubated for 30 minutes in the milk solution with a 1:10,000 dilution of Li-Cor near-infrared fluorescence secondary antibodies. The blots were scanned using Li-Cor’s Near-InfraRed fluorescence Odyssey CLx Imaging System, and quantifications were done using LiCor’s ImageStudio software complementary of Odyssey.

### Immunofluorescence

Immunofluorescence was conducted as previously described (Liang et al., 2018). Briefly, cells were fixed in 4% (wt/vol) paraformaldehyde for 15 min followed by permeabilization using 0.1% Triton-X100 in PBS for 10 min. Cells were then blocked in 1% BSA in PBS for 20 min. Primary antibodies were incubated for 1hr at room temperature, followed by three PBS washes for 5min each. Secondary antibodies were incubated for 30 min at room temperature, followed by three PBS washes for 5min each. Coverslips were mounted onto glass slides using ProLong Gold Antifade reagent with or without DAPI addition for nucleus visualization. Images were taken using either Zeiss LSM 710 Axio Observer (in Berkeley) or Leica TCS SP8 confocal microscope (in ETH Zurich) with 63x objective lens and post-processed in Adobe Photoshop for specific inset enlargement and RGB channel separation. Colocalization analysis in Figure 6J was determined by Pearson’s Correlation coefficient using ImageJ with colocalization plugin from McMaster Biophotonics Facility (MBF). The frequency scatterplot in Fig S5C was generated using the same plugin.

### Primary and secondary antibodies for Western blotting and immunofluorescence

All primary and secondary antibodies used were detailed in Supplemental table S6.

### MitoTracker

The MitoTracker assay was performed according the manufacturer’s protocol (ThermoFisher Catalog #M7512). Cells were plated 48 hours before starvation. Cell starvation and drug concentrations was performed according to protocols described above. The MitoTracker Red CMXRos was dissolved in DMSO for a stock concentration of 1 mM. MitoTracker was added to samples such that the concentration in each well was 50 nM. The cells were incubated for 30 minutes, washed with media, and then fixed in 4% formaldehyde. The cells were stained with calnexin according to the immunofluorescence protocol.

### ATP Assay

The assay was performed according to the manufacturer’s protocol for the CellTiter-Glo 2.0 reagent (Promega). Briefly, the cells treated with starvation were starved for 25 hours. The cells treated with rotenone or antimycin A were used as positive controls and cells were treated for 1 hour. Cells were harvested, washed with PBS, and counted and normalized. The cells were spun down again and resuspended such that there were 25,000 cells per 50 µL of PBS. 50 µL of PBS was added to each well in an opaque-walled 96-well plate. Each sample was done in technical triplicate. Wells with PBS, but no cells, were used as a blank control. 50 µL of CellTiter-Glo 2.0 reagent was added to each well. The plate was placed on an orbital shake for 2 minutes, followed by a 10 minute bench-top incubation to stabilize the signal. Sample luminescence was determined by the SpectraMax M2 Microplate Reader (Molecular Devices).

### Mitochondrial Respiration Measurements

Mitochondrial activity was determined using the Seahorse Flux Analyzer XF24 according to the manufacturer’s protocol. Briefly, 4 × 10^4^ HCT116 cells were seeded on XF24-well cell culture microplates. After 24hr, growth medium was exchanged with XF assay base medium supplemented with 1 mM sodium pyruvate, 2mM L-glutamine, and 10mM D-glucose (pH 7.4). The microplates were incubated at 37°C without CO_2_ for 1 hr prior to the assay. Samples were mixed for 3 min, time delayed for 2 min, and measured for 3 min. Oligomycin (1 µM), FCCP (1 µM), and rotenone / antimycin (0.5 µM) were sequentially injected at the indicated time points. OCR data were normalized by protein concentration and the average values were taken for each experiment. Seven replicates were performed for each cell line. The mean +/- SEM was determined and statistical significance was evaluated using the Student’s *t* test with a P value <0.05.

### Statistical analysis

All analysis was performed using data from three independent analysis, unless otherwise stated. Statistical analyses were performed in PRISM6 software using either paired Student’s *t*-test or ANOVA and are indicated in the figure legend.

---

**Figure S1:**
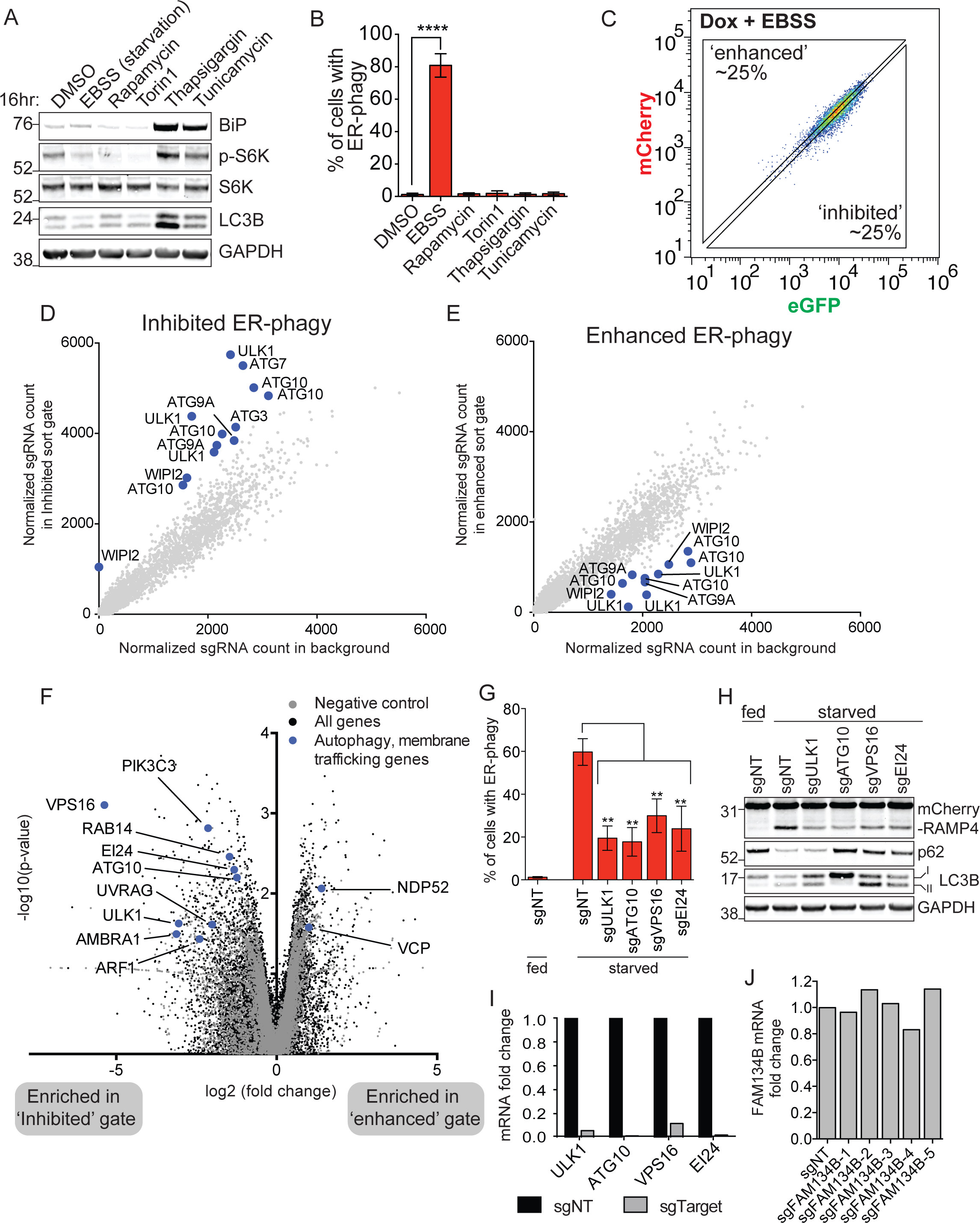
Related to Figure 1. A. EBSS (amino acid starvation), rapamycin, and Torin1 show a decrease in p-S6K while thapsigargin inhibits LC3B turnover. HCT116 EATR cells were treated with the indicated media or drugs (Rapamycin: 1µM, Torin1: 10µM, Thapsigargin: 1µM, Tunicamycin: 0.5µg/ml) for 16hr. Cells were then harvested for Western blotting to check for UPR and autophagy response.
B. Only starvation (using EBSS) inhibits ER-phagy. The same cells and treatment conditions in Fig S1A were also subjected to FACS measurement of ER-phagy. Data represents mean ± SD of three biological replicates. P value indicates two-tailed unpaired t-test (****, P< 0.0001).
C. Gating strategy for ER-phagy screen. HCT116 CRISPRi EATR cells transduced with the CRISPRi library of sgRNAs were starved for 16hr and subjected for FACS measurement for ER-phagy. Based on the EATR assay, the top and bottom 25% of cells correspond to sgRNA knockdown that results in ‘enhanced’ and ‘inhibited’ ER-phagy, respectively and were processed for next generation sequencing of sgRNA barcode.
D. Multiple autophagy genes (highlighted in blue) were enriched in the ‘inhibited’ ER-phagy sort gate. The normalized sgRNA count from the ‘Inhibited’ ER-phagy sort gate was plotted against the normalized sgRNA count of the background sample. The labelled genes are enriched in the ‘Inhibited’ ER-phagy gate upon knockdown and indicate active transcription start sites (TSS) that are being used in HCT116 cells.
E. Autophagy genes (highlighted in blue) that are depleted in the ‘enhanced’ ER-phagy sort gate corresponds to the genes that are enriched in the ‘inhibited’ sort gate of (D).
F. Autophagy and membrane trafficking genes are hits in the ER-phagy screen. Those genes that are significantly enriched in either the ‘inhibited’ or ‘enriched’ gate are indicated in blue. Volcano plot describes data from the genome-wide CRISPRi screen. All negative control sgRNAs are indicated in grey and targeted sgRNAs are indicated in black. Data were generated from two biological replicates. log2 fold change and Mann-Whitney P-value were calculated as described <u>(Horlbeck et al. 2016)</u>.
G. Knockdown of genes involved in general autophagy also inhibit ER-phagy. HCT116 CRISPRi EATR cells were transduced with sgRNAs targeting the indicated candidate genes reported from the CRISPRi genome wide screen. Cells were starved for 16hr and subjected to FACS measurement for ER-phagy. Data represents mean ± SD of three biological replicates. P value indicates two-tailed unpaired t-test (**, P< 0.01).
H. CCER assay complements the EATR data in Fig S1G. HCT116 CRISPRi CCER cells were also transduced with the same sgRNAs as Fig S1G. Cells were then starved for 16hr and harvested for Western blot analysis and immunoprobed for two general autophagy markers, p62 and LC3B.
I. The same cell lines in Fig S1H were harvested for qRT-PCR analysis to measure the knockdown efficiency of individual sgRNAs.
J. The FAM134B sgRNAs used in this CRISPRi screen do not effectively knock down FAM134B. HCT116-CRISPRi EATR cells stably expressing FAM134B sgRNAs were harvested for qRT-PCR to determine the knockdown efficiency.

**Figure S2:**
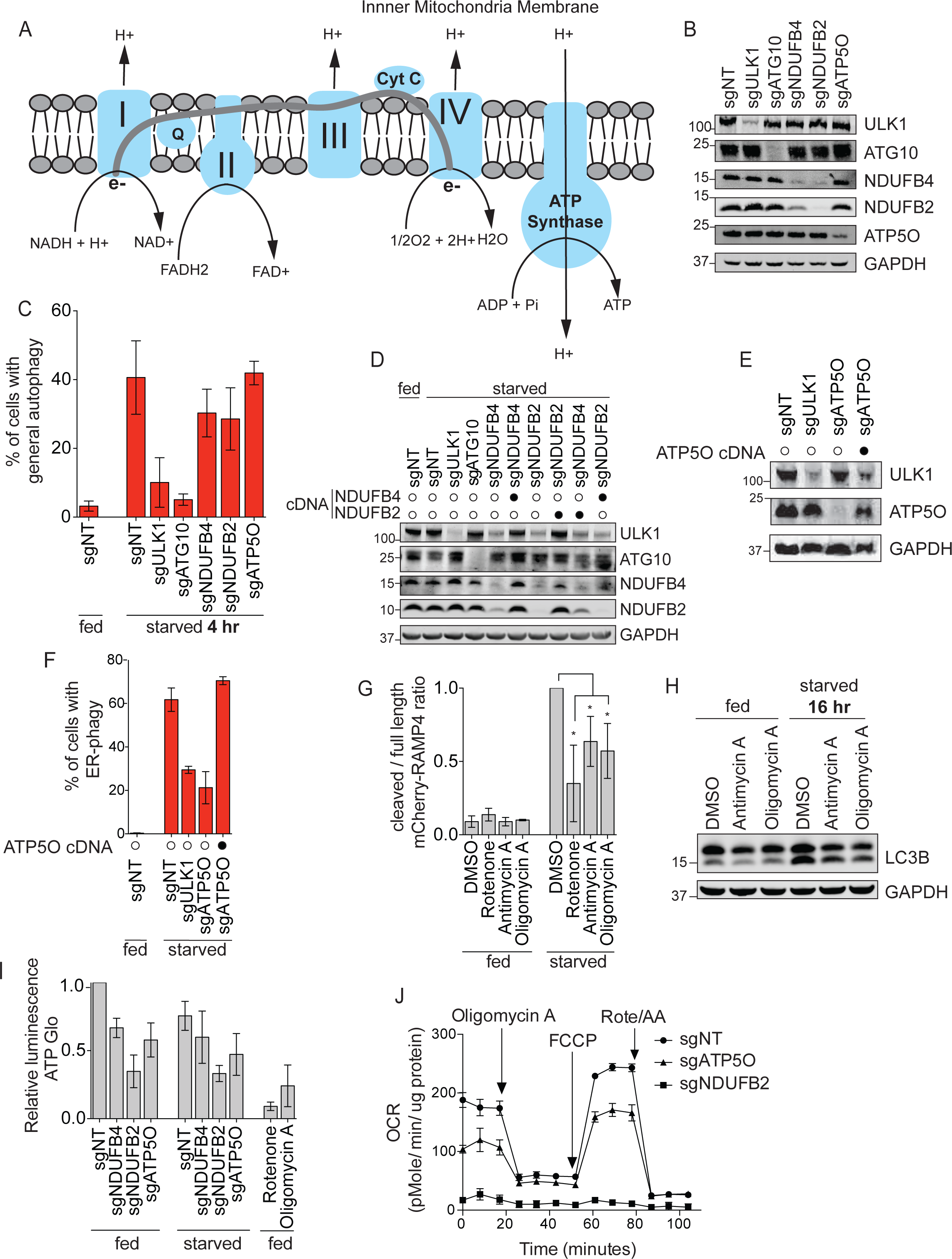
Related to Figure 2 and Figure 3. A. Schematic of the oxidative phosphorylation (OXPHOS) pathway.
B. Cells from Fig 2B were harvested to assess the knockdown efficiency of each sgRNA by Western blotting.
C. Knockdown of ULK1 and ATG10 inhibits general autophagy after 4 hours of starvation while NDUFB4, NDUFB2, and ATP5O does not. HCT116 CRISPRi cells expressing eGFP-mCherry-LC3B were transduced with sgRNAs targeting either ULK1, ATG10, NDUFB4, NDUFB2, or ATP5O. Cells were starved for 4hr before FACS measurement for general autophagy. Data represents mean ± SD of four biological replicates.
D. Cells from the same experiment as Fig 2D were harvested for Western blot analysis to verify the protein levels of ULK1, ATG10, NDUFB4 and NDUFB2.
E. Cells from the same experiment as Fig S2F were harvested for Western blot analysis to verify the protein levels of ULK1 and ATP5O.
F. Knockdown of ATP5O inhibits ER-phagy, but ATP5O cDNA rescues ER-phagy. HCT116 CRISPRi EATR cells were transduced with ATP5O cDNA constructs, and then transduced with sgRNAs targeting ULK1 or ATP5O. Cells were starved for 16hr before FACS measurement for ER-phagy. Data presented as mean ± SD of three biological replicates.
G. Rotenone, antimycin A, and oligomycin A significantly inhibit ER-phagy during starvation. Densitometry measurement of the ratio between the cleaved and full length mCherry-RAMP4 in Fig 3C. Data represents mean ± SD of three biological replicates. P value indicates two-tailed unpaired t-test (*, P < 0.05).
H. General autophagy is unaffected by antimycin A and oligomycin A treatment in starvation conditions. HCT116 cells were treated with the indicated small molecule inhibitors of antimycin A, or oligomycin A for 24hr and then were subsequently starved for 16hr with treatment of small molecule inhibitors. Cells were lysed for Western blotting and immunoprobed for the indicated proteins.
I. NDUFB2, NDUFB4 and ATP5O depletions affect cellular ATP levels. HCT116 CRISPRi cells were transduced with sgRNAs targeting either NDUFB4, NDUFB2, or ATP5O and starved for 16hr or treated with rotenone or oligomycin A. Cells were collected for ATP Glo luminescence assay. Data represents mean ± SD of three biological replicates.
J. Oxygen consumption is reduced in NDUFB4, NDUFB2, and ATP5O knockdown cells. HCT116 CRISPRi cells were transduced with sgRNAs targeting either NDUFB2 or ATP5O and the Seahorse Flux Analyzer was conducted to determine the oxygen consumption rate.

**Figure S3:**
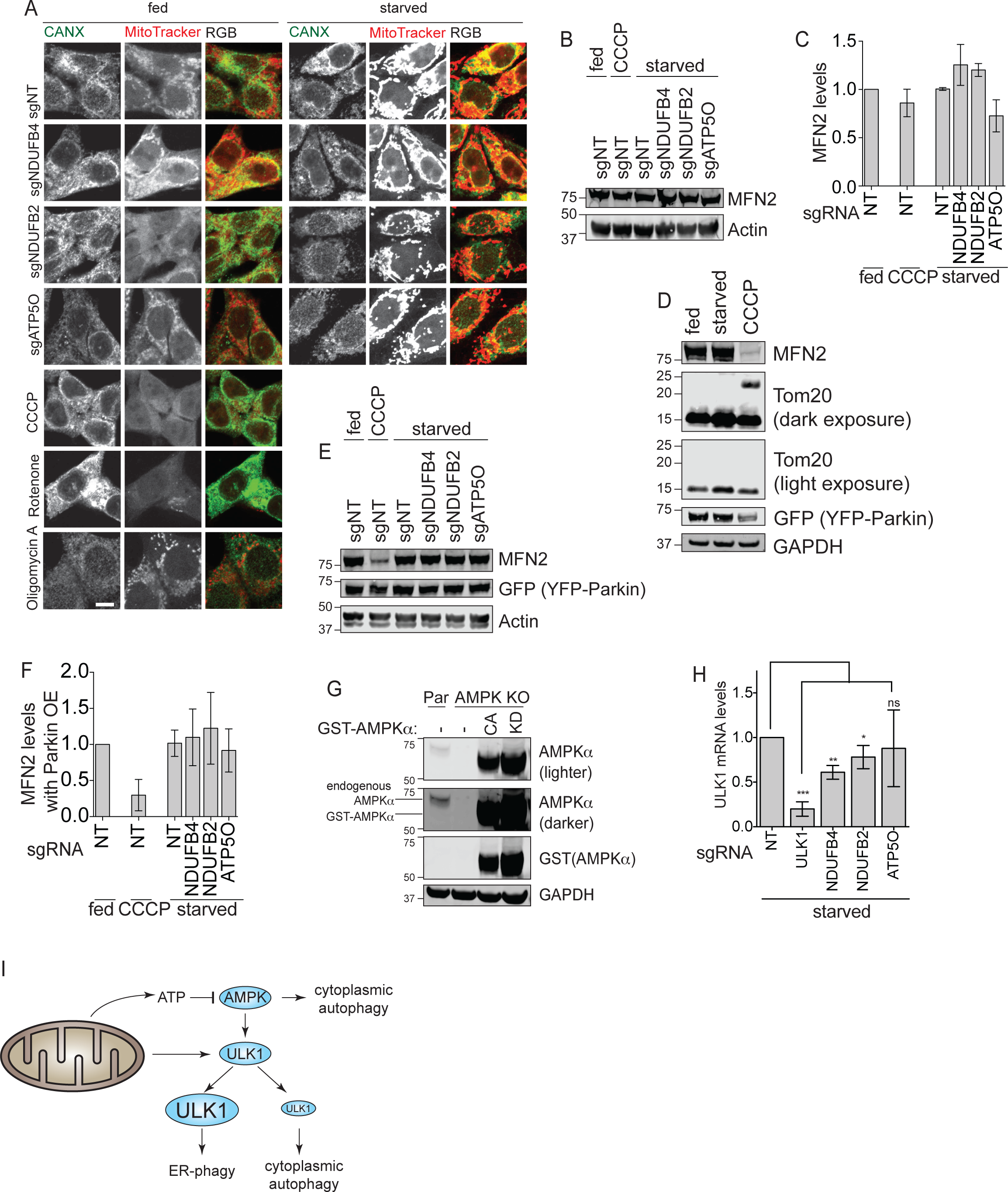
Related to Figure 4. A. sgNDUFB4, sgNDUFB2, and sgATP5O showed no major differences in MitoTracker staining compared to sgNT. HCT116 CRISPRi cells were transduced with sgRNAs targeting either NDUFB4, NDUFB2, or ATP5O or treated with indicated small molecule inhibitors. Cells were subsequently treated with MitoTracker, then fixed and immunostained for calnexin. Representative images are shown. Scale bar represents 10µm.
B. Disruption of the OXPHOS pathway does not induce mitophagy based on the protein levels of MFN2. HCT116 CRISPRi cells were transduced with sgRNAs targeting either NDUFB4, NDUFB2, or ATP5O and starved for 16hr or treated with CCCP. Cells were harvested for western blot analysis and immunoprobed for the indicated proteins. A representative western blot is shown.
C. Densitometry measurement of the ratio between MFN2/actin from Fig S3B. Data represents mean ± SD of two biological replicates.
D. HCT116 CRISPRi were transduced with YFP-Parkin construct and treated with starvation or CCCP (10 uM) for 16hr. Cells were harvested for Western blot analysis and immunoprobed for the indicated proteins to verify mitophagy in CCCP-treated cells.
E. MFN2 is not degraded in Parkin overexpression cell lines. HCT116 CRISPRi-YFP-Parkin cells were transduced with sgRNAs targeting either NDUFB4, NDUFB2, or ATP5O and starved for 16hr or treated with CCCP. Cells were harvested for Western blot analysis and immunoprobed for the indicated proteins. A representative western blot is shown.
F. MFN2 is not degraded in Parkin overexpression cell lines during OXPHOS disruption. Densitometry measurement of the ratio between MFN2/actin from Fig 3SE. Data represents mean ± SD of three biological replicates.
G. AMPKα knockout cell lines stably expressing either catalytically active (CA) or kinase dead (KD) AMPKα were generated to test the role of AMPK in ER-phagy. “Par” represents HCT116 CRISPRi EATR cells. AMPKα KO (knockout) cells were generated using CRISPR/Cas9 (sgRNA sequences and methods are described in the methods section). GST-AMPKα-CA (catalytically active) was re-expressed in the AMPKα KO background. Note that the catalytically active AMPKα is a truncation form of AMPKα, resulting in the smaller size. GST-AMPKα-KD (kinase dead) is derived from GST-AMPKα-CA with a K47R point mutation.
H. ULK1 mRNA levels are significantly reduced during starvation when NDUFB4 and NDUFB2 are knocked down. HCT116 CRISPRi EATR cells were transduced with sgRNAs targeting ULK1, NDUFB4, NDUFB2, or ATP5O and starved for 16hr. RNA was extracted and cDNA was synthesized for qRT-PCR. Data presented as mean ± SD of three biological replicates. P value indicates two-tailed paired t-test (*, P < 0.05, **, P < 0.01, ***, P < 0.001).
I. Schematic representation of the interplay between mitochondrial metabolism, ULK1, and ER-phagy. Disruption of the mitochondrial OXPHOS pathway results in decreased ATP levels and leads to activation of cytoplasmic autophagy. In parallel, impairment of the mitochondrial OXPHOS also transcriptionally lowers ULK1 levels to an extent that inhibits ER-phagy while still permits cytoplasmic autophagy to take place.

**Figure S4:**
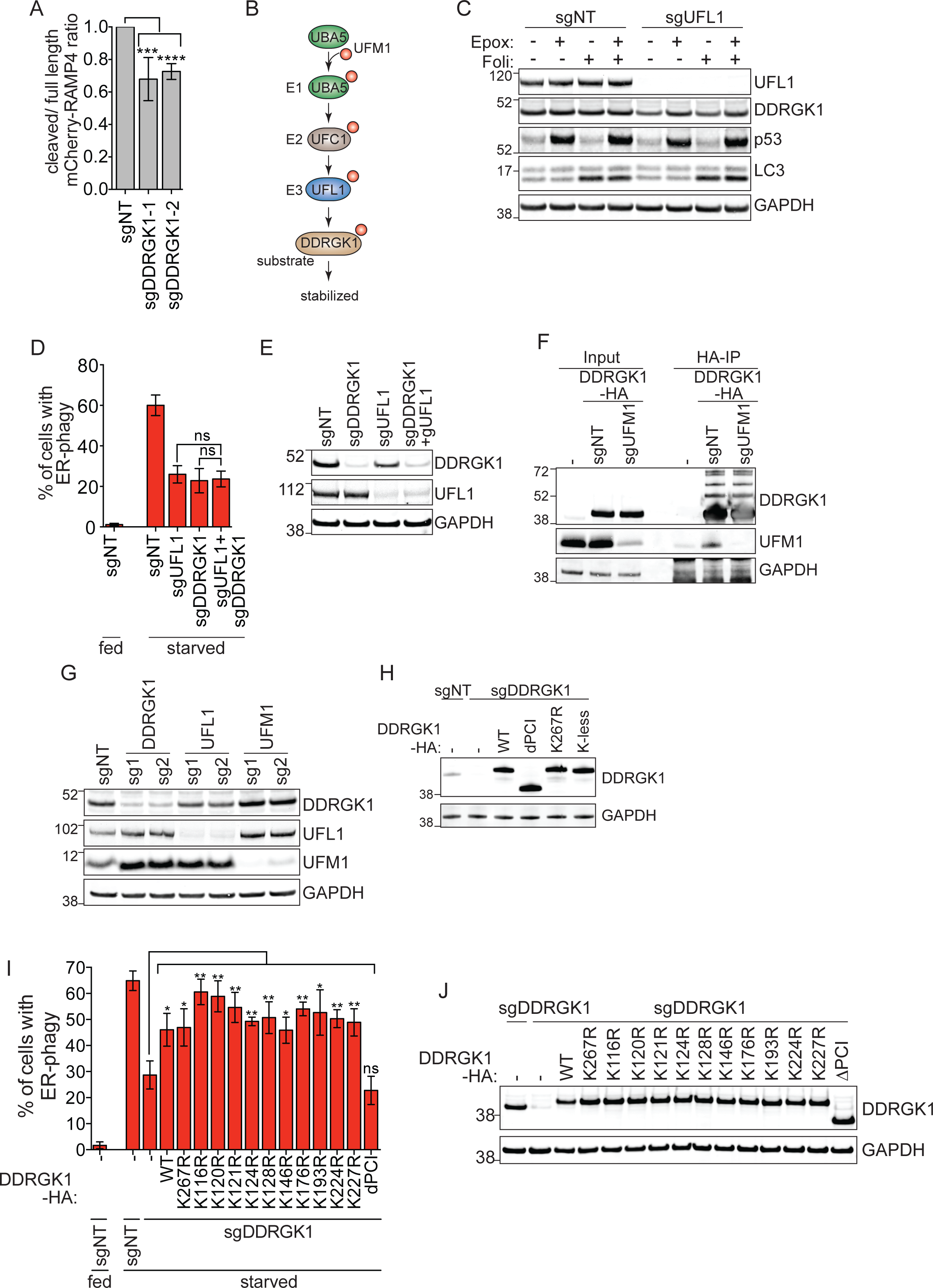
Related to Figure 5 and 6. A. Densitometry measurement of the ratio between the cleaved and full length mCherry-RAMP4 in Fig 5C. Data represents mean ± SD of five biological replicates. P value indicates two-tailed unpaired t-test (***, P < 0.001, ****, P<0.0001).
B. Schematic illustration of the three-step enzymatic reaction of the UFMylation cascade. UBA5 acts as an E1 enzyme to activate UFM1 and UFC1 acts as an E2 conjugating enzyme that interacts with the E3 ligase, UFL1. UFL1 recognizes and transfer UFM1 from UFC1 to its target substrate. In this case, DDRGK1 is reported as a substrate of UFMylation.
C. UFL1 protein expression prevents DDRGK1 degradation via the proteasomal degradation pathway. HCT116 CRISPRi cells were transduced with sgRNA targeting UFL1. Cells were treated with Epoxomicin (100nM) or Folimycin (100nM) for 6hr and then harvested for Western blot analysis. p53 and LC3B were used as positive controls for proteasomal and lysosomal inhibitions, respectively.
D. UFL1 and DDRGK1 act in series during ER-phagy. HCT116 CRISPRi EATR cells were transduced with the indicated sgRNAs and then starved for 16hr before FACS analysis for ER-phagy. Data represents mean ± SD of three biological replicates. Statistical significance was determined based on two-tailed unpaired t-test.
E. The same cell lines in Fig S4B were harvested for Western blotting to verify knockdown of targeted genes.
F. Knockdown of UFM1 does not prevent the appearance of higher molecular weight species of DDRGK1. HCT116 CRISPRi cells stably expressing DDRGK1-HA construct were transduced with sgRNA targeting UFM1. Cells were then harvested for HA-immunoprecipitation and Western blotted for the indicated proteins.
G. Knockdown of UFL1 but not UFM1 reduces DDRGK1 protein levels. HCT116 CRISPRi EATR cells used in Fig 6C were harvested for Western blotting to assess the protein levels of DDRGK1 upon knockdown of the targeted genes.
H. The cDNA of DDRGK1-HA mutant variants in Fig 6F were stably expressed in HCT116 cells to verify their respective protein sizes in DDRGK1 knockdown cells.
I. All individual Lysine mutant constructs of DDRGK1 are able to rescue ER-phagy in DDRGK1 knockdown cells. HCT116 CRISPRi EATR cells with sgDDRGK1 were transduced with the indicated DDRGK1-HA mutant constructs. Cells were then starved for 16hr before FACS measurement of ER-phagy. Data represents mean ± SD of three biological replicates. P value indicates two-tailed unpaired t-est. (*, P<0.05, **, P< 0.01).
J. The cell lines generated for Fig S4I were harvested for Western blotting to verify the expression of the indicated DDRGK1-HA mutant constructs.

**Figure S5:**
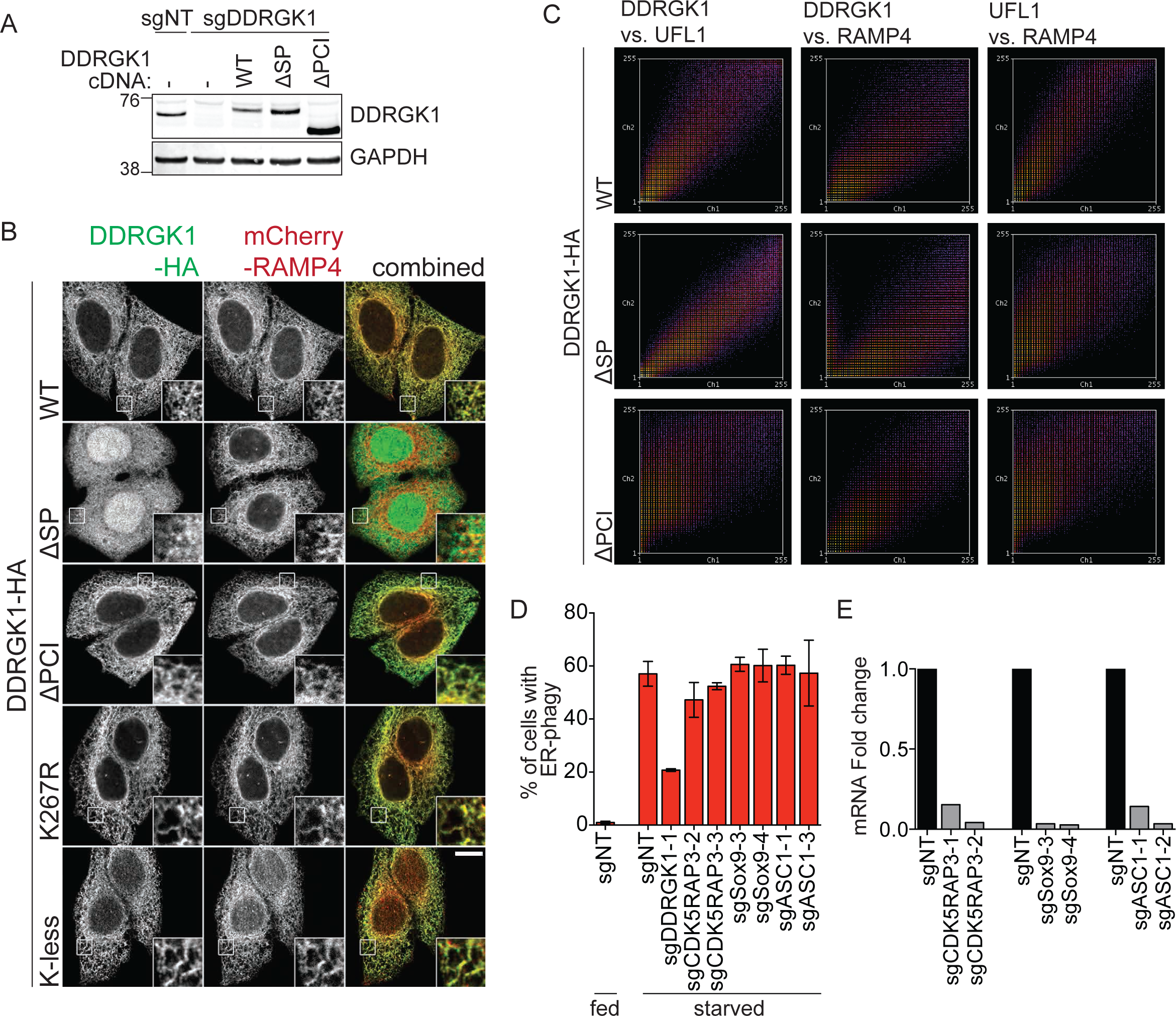
Related to Figure 6. A. The cell lines generated for Fig 6K were harvested for Western blotting to verify the expression of the indicated DDRGK1-HA mutant constructs.
B. The ER signaling peptide (SP) of DDRGK1 is required for its localization to the ER. HeLa cells stably expressing mCherry-RAMP4 were transduced with the indicated DDRGK1-HA constructs. Cells were then fixed and stained for HA-epitope. Insets represent three-fold enlargement of the boxed areas. Scale bar represents 10µm.
C. Additional colocalization data analysis of Fig 6J for DDRGK1, UFL1 and the ER marker, RAMP4 based on frequency of colocalization. Conditions with good or near perfect colocalization have good correlation between the X- and Y-axes. Representative frequency scatterplot from each condition is presented.
D. None of the previously reported targets of UFMylation are involved in ER-phagy regulation. HCT116 CRISPRi EATR cells were transduced with sgRNAs targeting CDK5RAP3, Sox9 or ASC1 and starved for 16hr before FACS measurement of ER-phagy. Data represents mean ± SD of three biological replicates.
E. The same cell lines used in Fig S5D were harvested for qRT-PCR to assess the knockdown efficiency of each sgRNA.

**Figure S6:**
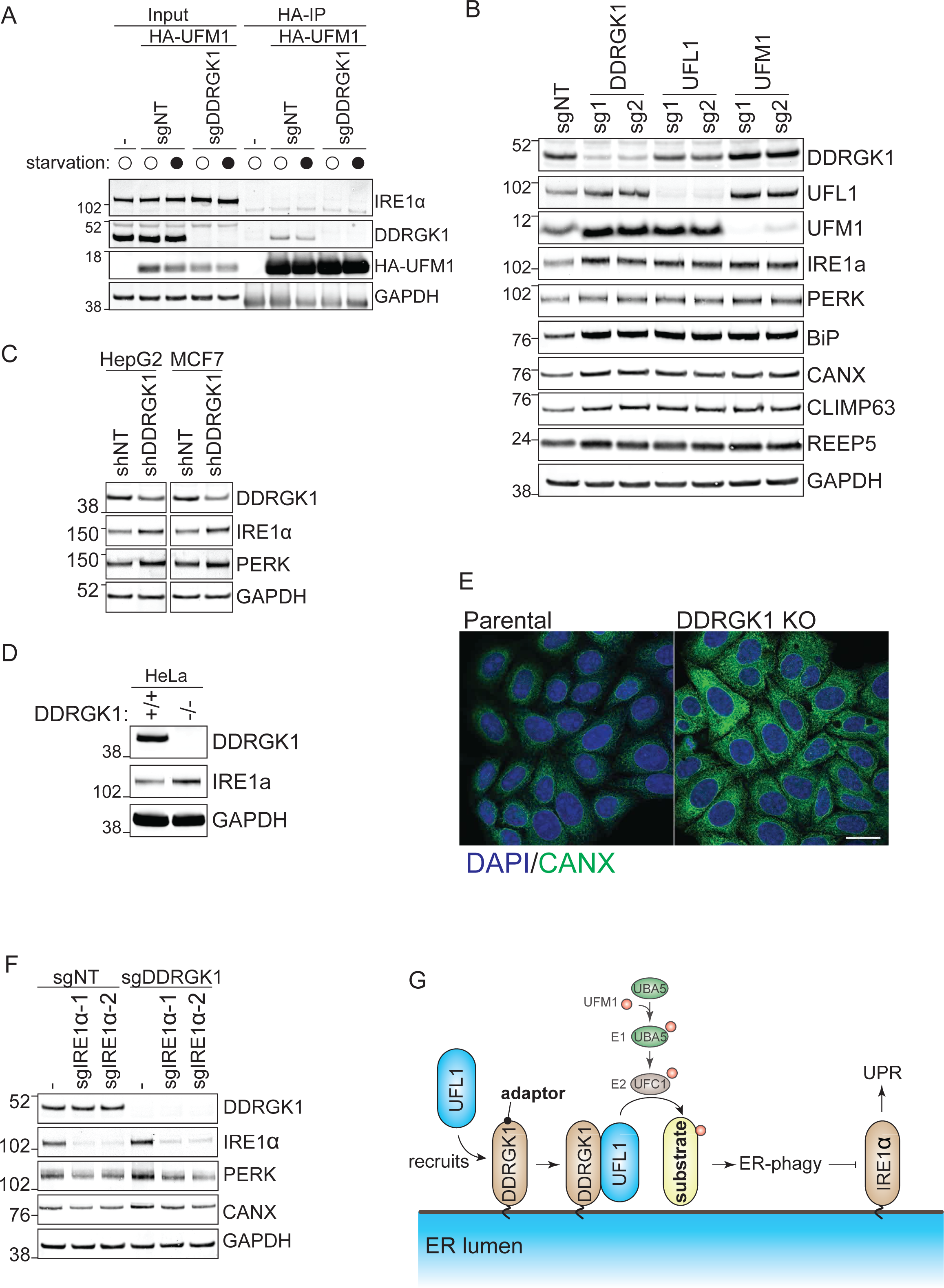
Related to Figure 7. A. IRE1α is not a target of DDRGK1-mediated UFMylation. HCT116 CRISPRi cells stably expressing HA-UFM1 were transduced with sgRNA targeting DDRGK1. Cells were then starved for 16hr before being harvested for HA- immunoprecipitation of UFM1 and Western blot analysis of its interaction with IRE1α and DDRGK1.
B. The same experiment shown in Fig S4G were further probed for the indicated ER or UPR markers. The densitometry measurements for IRE1α, BiP, CANX, CLIMP63 and REEP5 are shown in biological triplicates in Fig 7A.
C. DDRGK1 depletion results in the upregulation of UPR markers in various cell lines. HepG2 and MCF7 cells were transduced with shRNAs targeting DDRGK1. Cells were then harvested for Western blot analysis of DDRGK1 knockdown efficiency and UPR response.
D. HeLa cells were transduced with shRNAs targeting DDRGK1. Cells were then harvested for Western blot analysis of DDRGK1 knockdown efficiency and UPR response.
E. The same HeLa cell lines generated in S6D were fixed and stained for an ER marker (CANX). Scale bar represents 20µm.
F. IRE1α depletion prevents UPR response in DDRGK1 knockdown cells. The same cell lines in Fig 7C were harvested for Western blotting to measure the protein levels of the indicated UPR genes.
G. Proposed model for UFMylation regulation on ER-phagy. DDRGK1 acts as an adaptor/anchor for UFL1 to recruit the latter to the ER surface. ER-localized DDRGK1-UFL1 then UFMylates a yet-to-be-identified substrate(s) that regulates ER-phagy. Loss of UFMylation factors results in ER-phagy inhibition and accumulation of ER stress. This in turn triggers UPR via the IRE1α pathway.

**Table S1.**
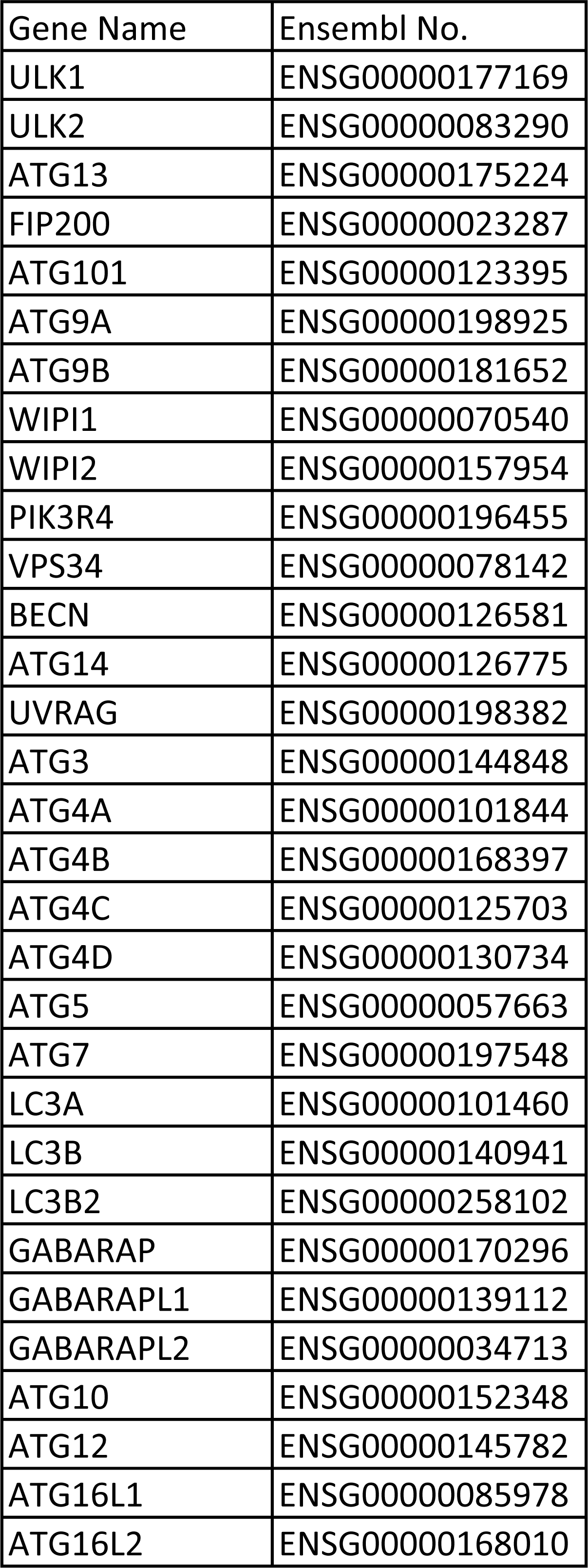
General ATG targets.

**Table S2.**
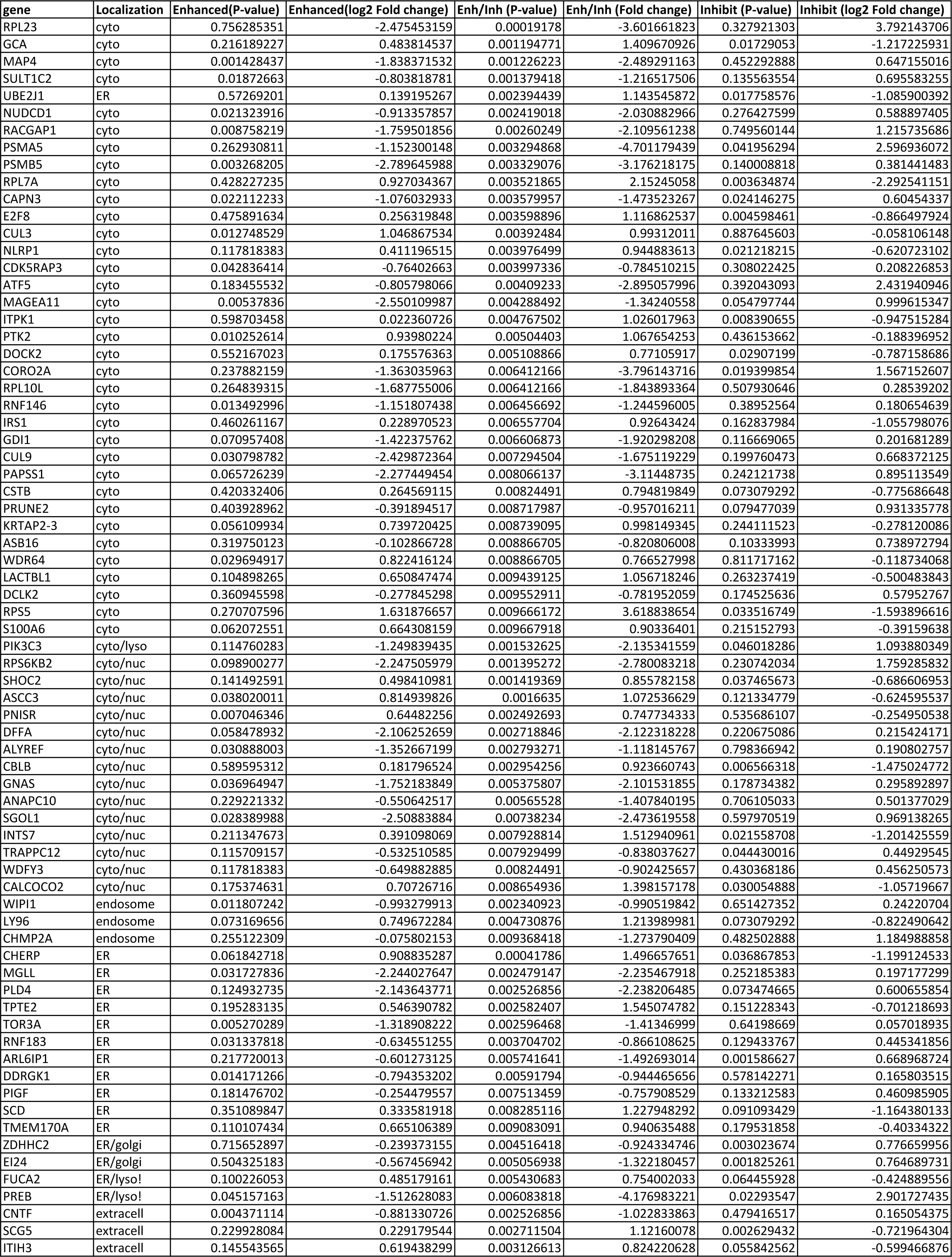
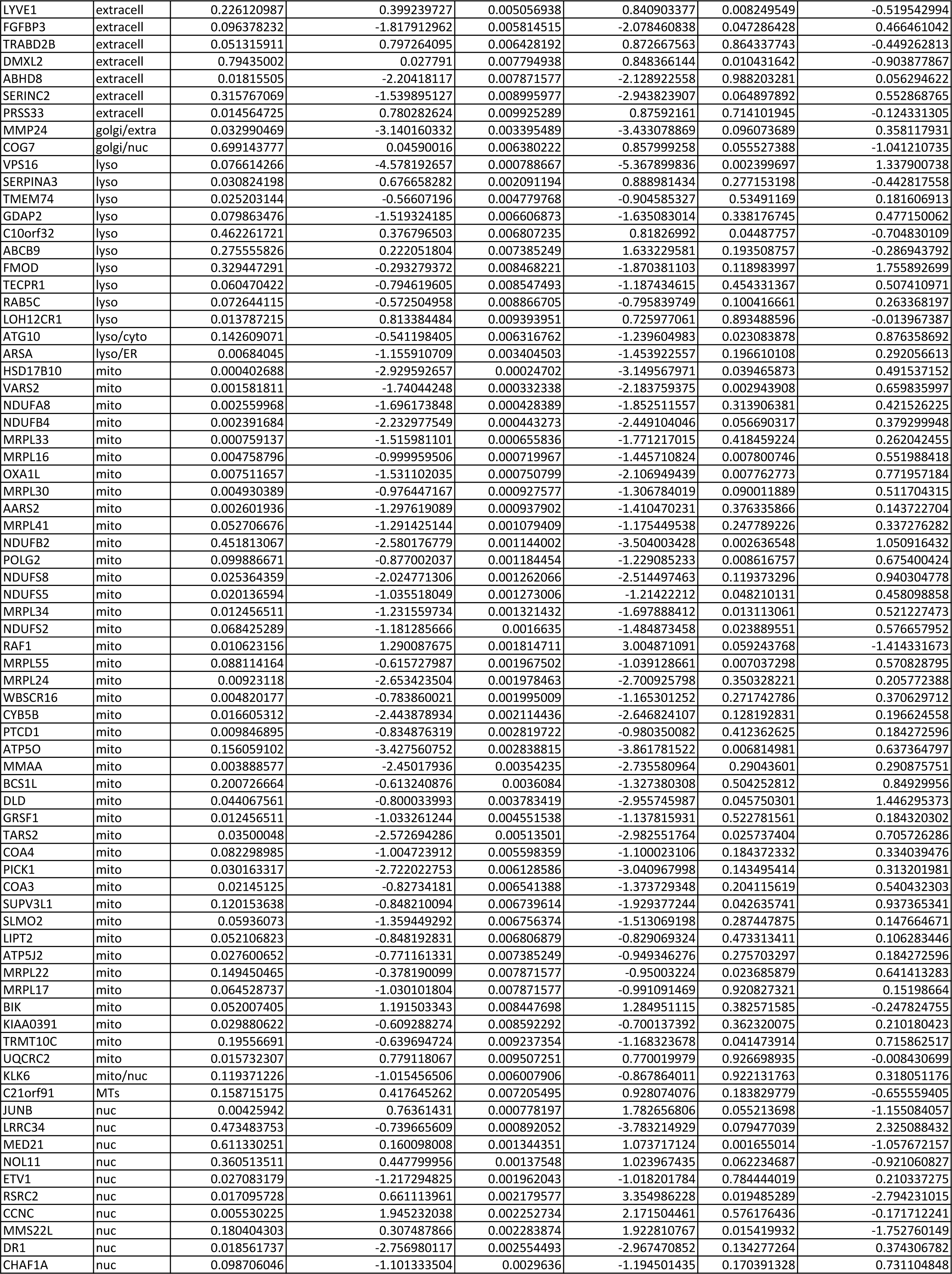
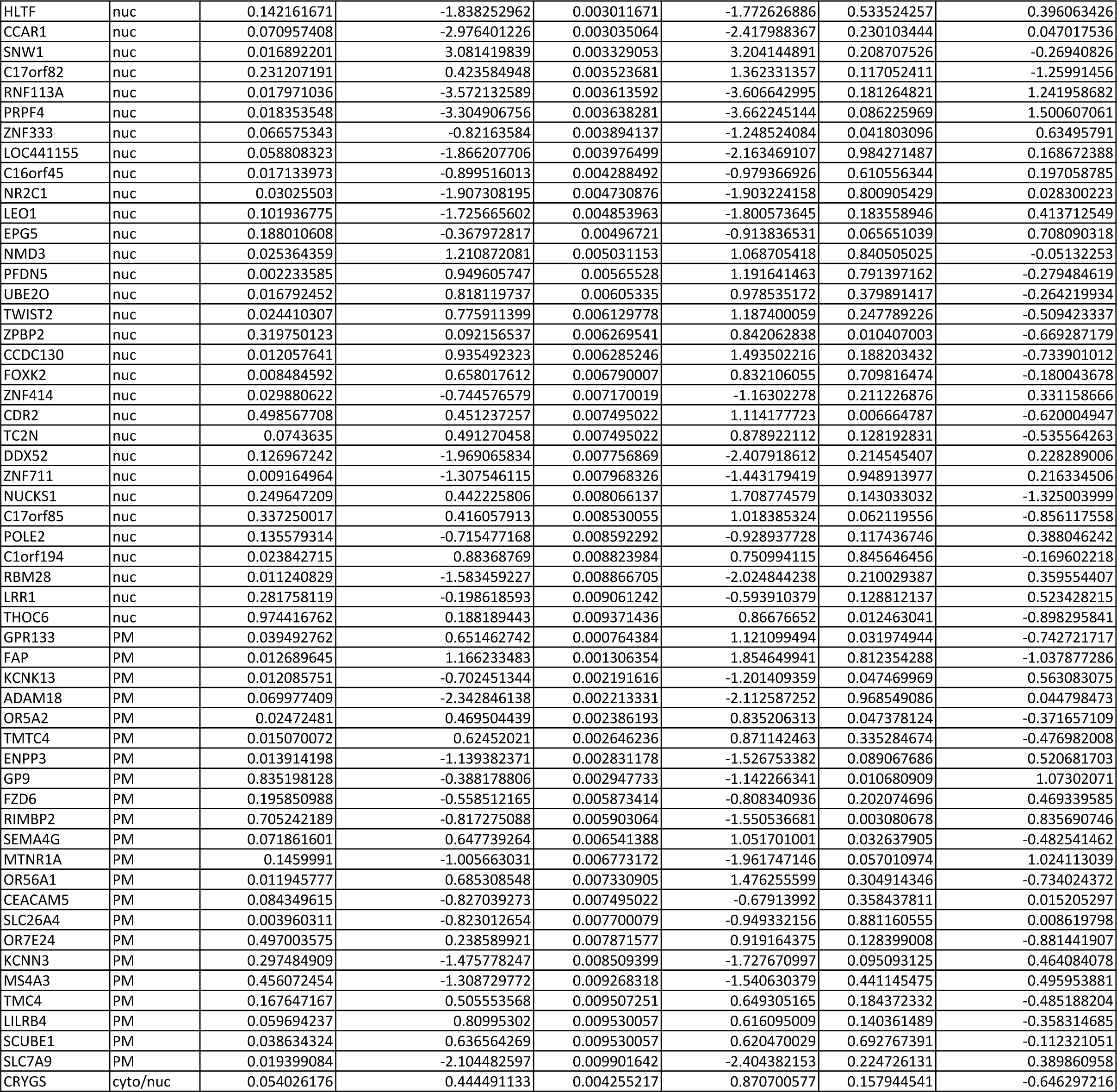
high confidence hits with Enhanced/Inhibition P<0.01.

**Table S3.**
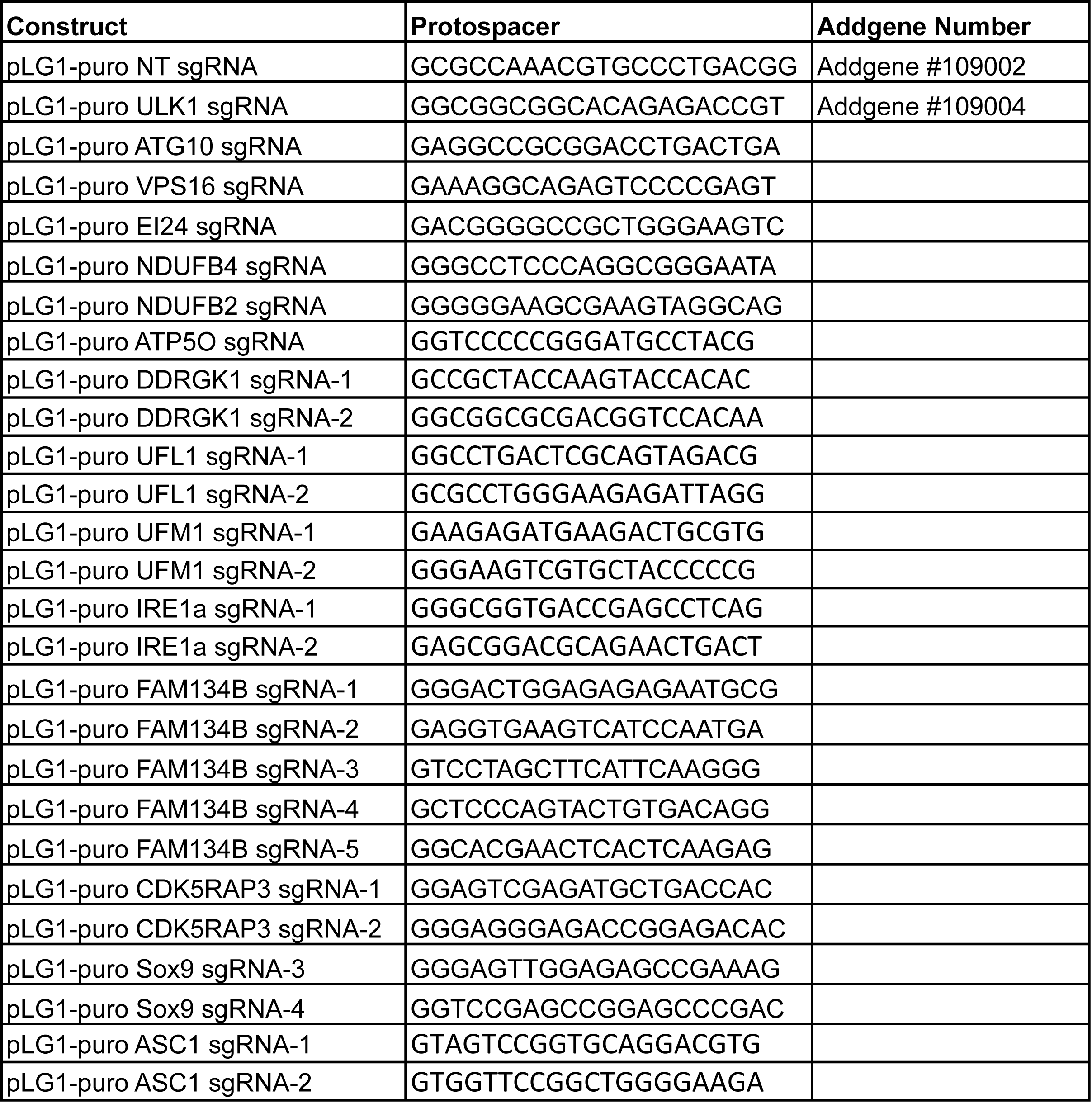
sgRNA constructs.

**Table S4.** Overexpression constructs.

| **Plasmids** | **ORF source** |
| --- | --- |
| pLenti-X1-Neo-NDUFB4 | HCT116 cDNA |
| pLenti-X1-Neo-NDUFB2 | HCT116 cDNA |
| pLenti-X1-Neo-ATP5O | HCT116 cDNA |
| pLentiXI-Neo-GST-Constitutively Active AMPK | Addgene 27632 |
| pLentiXI-Neo-GST-Constitutively Active AMPK- Kinase Dead (K to R) | Addgene 27632 |
| pBMN-YFP-Parkin | Addgene 59416 |
| pLenti-X1-Neo-HA-hULK1 | Addgene 31963 |
| pLenti-X1-Neo-DDRGK1-WT-HA | HCT116 cDNA; NM_023935.2 |
| pLenti-X1-Neo-DDRGK1-dSP-HA | subcloned from pLenti-X1-Neo-DDRGK1-WT-HA |
| pLenti-X1-Neo-DDRGK1-dPCI-HA | subcloned from pLenti-X1-Neo-DDRGK1-WT-HA |
| pLenti-X1-Neo-DDRGK1-K267R-HA | subcloned from pLenti-X1-Neo-DDRGK1-WT-HA |
| pLenti-X1-Neo-DDRGK1-K-less-HA | subcloned from pLenti-X1-Neo-DDRGK1-WT-HA |
| pLenti-X1-Neo-DDRGK1-K116R-HA | subcloned from pLenti-X1-Neo-DDRGK1-WT-HA |
| pLenti-X1-Neo-DDRGK1-K120R-HA | subcloned from pLenti-X1-Neo-DDRGK1-WT-HA |
| pLenti-X1-Neo-DDRGK1-K121R-HA | subcloned from pLenti-X1-Neo-DDRGK1-WT-HA |
| pLenti-X1-Neo-DDRGK1-K124R-HA | subcloned from pLenti-X1-Neo-DDRGK1-WT-HA |
| pLenti-X1-Neo-DDRGK1-K128R-HA | subcloned from pLenti-X1-Neo-DDRGK1-WT-HA |
| pLenti-X1-Neo-DDRGK1-K146R-HA | subcloned from pLenti-X1-Neo-DDRGK1-WT-HA |
| pLenti-X1-Neo-DDRGK1-K176R-HA | subcloned from pLenti-X1-Neo-DDRGK1-WT-HA |
| pLenti-X1-Neo-DDRGK1-K193R-HA | subcloned from pLenti-X1-Neo-DDRGK1-WT-HA |
| pLenti-X1-Neo-DDRGK1-K224R-HA | subcloned from pLenti-X1-Neo-DDRGK1-WT-HA |
| pLenti-X1-Neo-DDRGK1-K227R-HA | subcloned from pLenti-X1-Neo-DDRGK1-WT-HA |
| pLenti-X1-Neo-HA-UFL1 | HCT116 cDNA; NM_015323.4 |
| pLenti-X1-Neo-IRE1A-no-tag | HCT116 cDNA; NM_001433.4 |
| pLenti-X1-Neo-IRE1A-K121Y-no-tag | subcloned from pLenti-X1-Neo-IRE1A-HA |
| pLenti-X1-Neo-HA-UFM1 | HCT116 cDNA; NM_016617.4 |
| pLKO.1-puro-shNon-targeting | Addgene 109012 |
| pLKO.1-puro-shDDRGK1 |  |

**Table S5.**
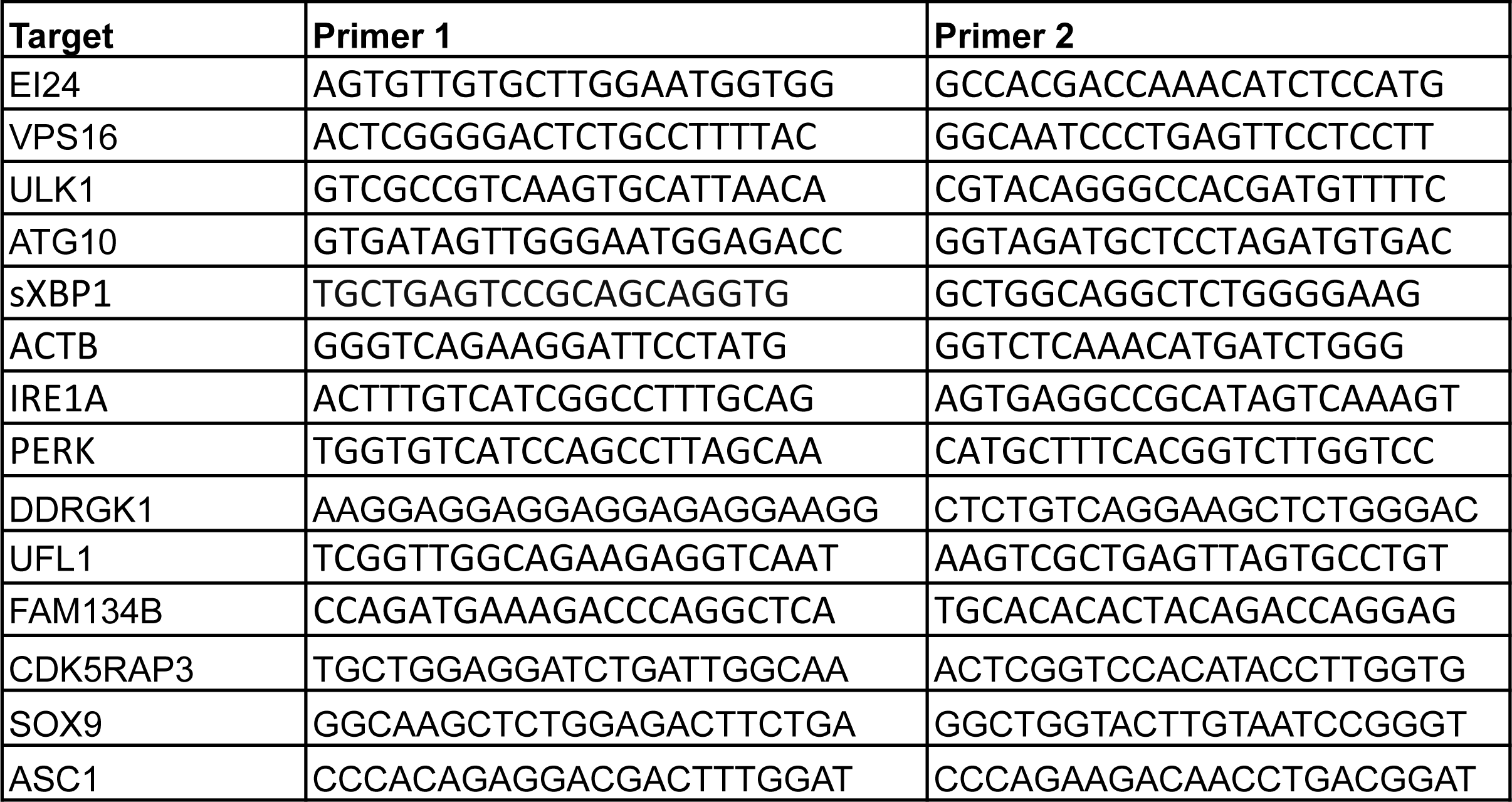
qRT-PCR primers.

**Table S6.**
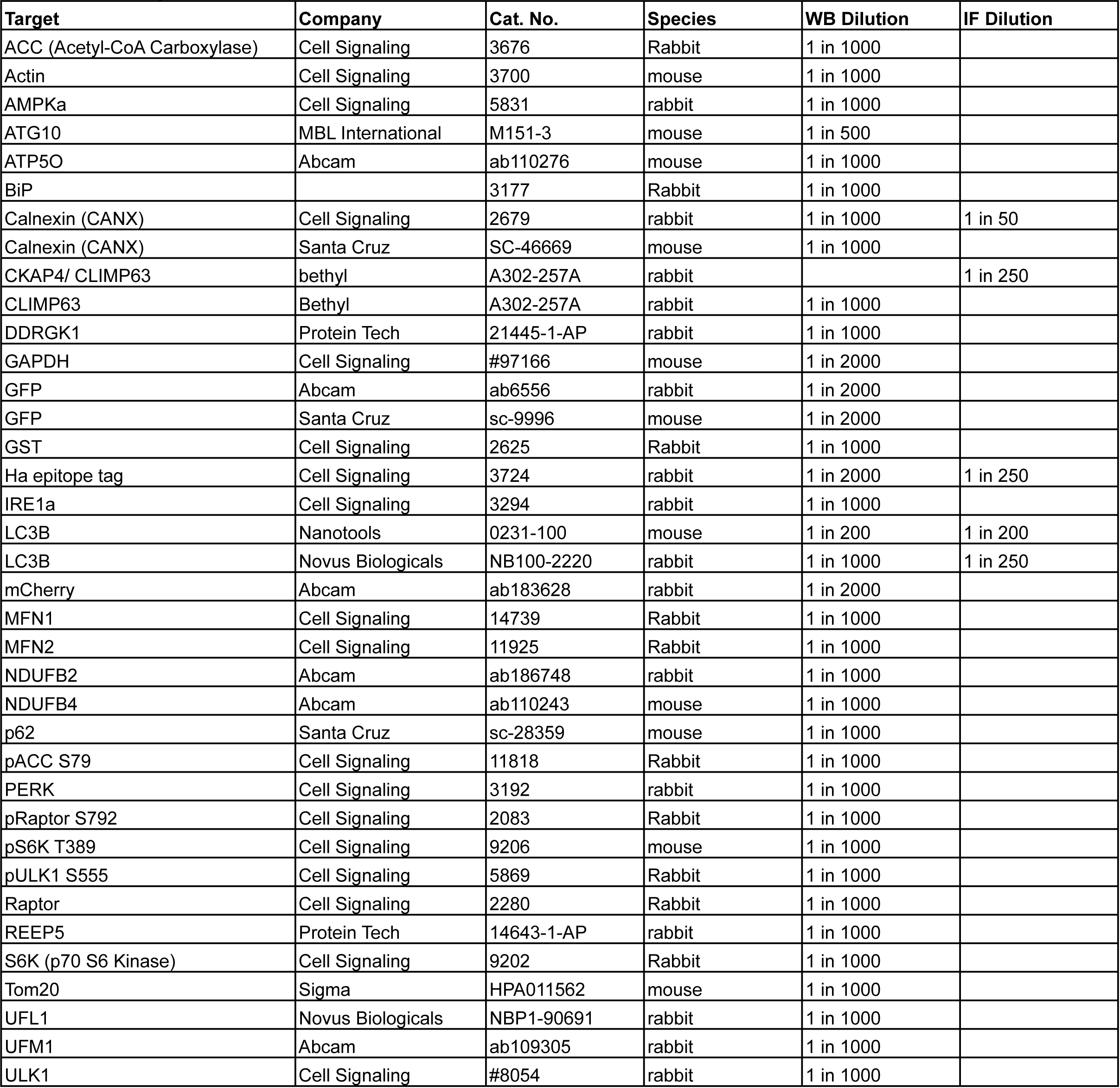
Antibody List.

## Bibliography

Abel, A., Fonknechten, N., Hofer, A., Dürr, A., Cruaud, C., Voit, T., Weissenbach, J., Brice, A., Klimpe, S., Auburger, G., et al. (2004). Early onset autosomal dominant spastic paraplegia caused by novel mutations in SPG3A. Neurogenetics 5, 239–243.

Adamson, B., Norman, T.M., Jost, M., Cho, M.Y., Nuñez, J.K., Chen, Y., Villalta, J.E., Gilbert, L.A., Horlbeck, M.A., Hein, M.Y., et al. (2016). A Multiplexed Single-Cell CRISPR Screening Platform Enables Systematic Dissection of the Unfolded Protein Response. Cell 167, 1867–1882.e21.

Amberger, J.S., Bocchini, C.A., Schiettecatte, F., Scott, A.F., and Hamosh, A. (2015). OMIM.org: Online Mendelian Inheritance in Man (OMIM®), an online catalog of human genes and genetic disorders. Nucleic Acids Res. 43, D789–98.

Binder, J.X., Pletscher-Frankild, S., Tsafou, K., Stolte, C., O’Donoghue, S.I., Schneider, R., and Jensen, L.J. (2014). COMPARTMENTS: unification and visualization of protein subcellular localization evidence. Database (Oxford) 2014, bau012.

Cai, Y., Pi, W., Sivaprakasam, S., Zhu, X., Zhang, M., Chen, J., Makala, L., Lu, C., Wu, J., Teng, Y., et al. (2015). UFBP1, a Key Component of the Ufm1 Conjugation System, Is Essential for Ufmylation-Mediated Regulation of Erythroid Development. PLoS Genet. 11, e1005643.

Chan, N.C., Salazar, A.M., Pham, A.H., Sweredoski, M.J., Kolawa, N.J., Graham, R.L.J., Hess, S., and Chan, D.C. (2011). Broad activation of the ubiquitin-proteasome system by Parkin is critical for mitophagy. Hum. Mol. Genet. 20, 1726–1737.

Chatr-Aryamontri, A., Oughtred, R., Boucher, L., Rust, J., Chang, C., Kolas, N.K., O’Donnell, L., Oster, S., Theesfeld, C., Sellam, A., et al. (2017). The BioGRID interaction database: 2017 update. Nucleic Acids Res. 45, D369–D379.

Daida, A., Hamano, S.-I., Ikemoto, S., Matsuura, R., Nakashima, M., Matsumoto, N., and Kato, M. (2018). Biallelic loss-of-function UBA5 mutations in a patient with intractable West syndrome and profound failure to thrive. Epileptic Disord. 20, 313–318.

Daniel, J., and Liebau, E. (2014). The ufm1 cascade. Cells 3, 627–638.

Deas, E., Wood, N.W., and Plun-Favreau, H. (2011). Mitophagy and Parkinson’s disease: the PINK1-parkin link. Biochim. Biophys. Acta 1813, 623–633.

DeJesus, R., Moretti, F., McAllister, G., Wang, Z., Bergman, P., Liu, S., Frias, E., Alford, J., Reece-Hoyes, J.S., Lindeman, A., et al. (2016). Functional CRISPR screening identifies the ufmylation pathway as a regulator of SQSTM1/p62. Elife 5.

Dodson, M.W., and Guo, M. (2007). Pink1, Parkin, DJ-1 and mitochondrial dysfunction in Parkinson’s disease. Curr. Opin. Neurobiol. 17, 331–337.

Egan, D.F., Shackelford, D.B., Mihaylova, M.M., Gelino, S., Kohnz, R.A., Mair, W., Vasquez, D.S., Joshi, A., Gwinn, D.M., Taylor, R., et al. (2011). Phosphorylation of ULK1 (hATG1) by AMP-activated protein kinase connects energy sensing to mitophagy. Science 331, 456–461.

Egunsola, A.T., Bae, Y., Jiang, M.-M., Liu, D.S., Chen-Evenson, Y., Bertin, T., Chen, S., Lu, J.T., Nevarez, L., Magal, N., et al. (2017). Loss of DDRGK1 modulates SOX9 ubiquitination in spondyloepimetaphyseal dysplasia. J. Clin. Invest. 127, 1475–1484.

Elangovan, M., Chong, H.K., Park, J.H., Yeo, E.J., and Yoo, Y.J. (2017). The role of ubiquitin-conjugating enzyme Ube2j1 phosphorylation and its degradation by proteasome during endoplasmic stress recovery. J. Cell Commun. Signal. 11, 265–273.

Fregno, I., Fasana, E., Bergmann, T.J., Raimondi, A., Loi, M., Soldà, T., Galli, C., D’Antuono, R., Morone, D., Danieli, A., et al. (2018). ER-to-lysosome-associated degradation of proteasome-resistant ATZ polymers occurs via receptor-mediated vesicular transport. EMBO J. 37.

Friedman, J.R., Lackner, L.L., West, M., DiBenedetto, J.R., Nunnari, J., and Voeltz, G.K. (2011). ER tubules mark sites of mitochondrial division. Science 334, 358–362.

Fumagalli, F., Noack, J., Bergmann, T.J., Cebollero, E., Pisoni, G.B., Fasana, E., Fregno, I., Galli, C., Loi, M., Soldà, T., et al. (2016). Translocon component Sec62 acts in endoplasmic reticulum turnover during stress recovery. Nat. Cell Biol. 18, 1173–1184.

Geisler, S., Holmström, K.M., Treis, A., Skujat, D., Weber, S.S., Fiesel, F.C., Kahle, P.J., and Springer, W. (2010). The PINK1/Parkin-mediated mitophagy is compromised by PD-associated mutations. Autophagy 6, 871–878.

Genschik, P., Sumara, I., and Lechner, E. (2013). The emerging family of CULLIN3-RING ubiquitin ligases (CRL3s): cellular functions and disease implications. EMBO J. 32, 2307–2320.

Gibson, D.G., Young, L., Chuang, R.-Y., Venter, J.C., Hutchison, C.A., and Smith, H.O. (2009). Enzymatic assembly of DNA molecules up to several hundred kilobases. Nat. Methods 6, 343–345.

Gilbert, L.A., Horlbeck, M.A., Adamson, B., Villalta, J.E., Chen, Y., Whitehead, E.H., Guimaraes, C., Panning, B., Ploegh, H.L., Bassik, M.C., et al. (2014). Genome-scale CRISPR-mediated control of gene repression and activation. Cell 159, 647–661.

Glauser, L., Sonnay, S., Stafa, K., and Moore, D.J. (2011). Parkin promotes the ubiquitination and degradation of the mitochondrial fusion factor mitofusin 1. J. Neurochem. 118, 636–645.

Grumati, P., Morozzi, G., Hölper, S., Mari, M., Harwardt, M.-L.I., Yan, R., Müller, S., Reggiori, F., Heilemann, M., and Dikic, I. (2017). Full length RTN3 regulates turnover of tubular endoplasmic reticulum via selective autophagy. Elife 6.

Guelly, C., Zhu, P.-P., Leonardis, L., Papić, L., Zidar, J., Schabhüttl, M., Strohmaier, H., Weis, J., Strom, T.M., Baets, J., et al. (2011). Targeted high-throughput sequencing identifies mutations in atlastin-1 as a cause of hereditary sensory neuropathy type I. Am. J. Hum. Genet. 88, 99–105.

Hamasaki, M., Noda, T., Baba, M., and Ohsumi, Y. (2005). Starvation triggers the delivery of the endoplasmic reticulum to the vacuole via autophagy in yeast. Traffic 6, 56–65.

Herzig, S., and Shaw, R.J. (2018). AMPK: guardian of metabolism and mitochondrial homeostasis. Nat. Rev. Mol. Cell Biol. 19, 121–135.

Horlbeck, M.A., Gilbert, L.A., Villalta, J.E., Adamson, B., Pak, R.A., Chen, Y., Fields, A.P., Park, C.Y., Corn, J.E., Kampmann, M., et al. (2016). Compact and highly active next-generation libraries for CRISPR-mediated gene repression and activation. Elife 5.

Huang, D.W., Sherman, B.T., and Lempicki, R.A. (2009). Systematic and integrative analysis of large gene lists using DAVID bioinformatics resources. Nat. Protoc. 4, 44–57.

Johnson, M.A., Vidoni, S., Durigon, R., Pearce, S.F., Rorbach, J., He, J., Brea-Calvo, G., Minczuk, M., Reyes, A., Holt, I.J., et al. (2014). Amino acid starvation has opposite effects on mitochondrial and cytosolic protein synthesis. PLoS ONE 9, e93597.

Karbowski, M., and Youle, R.J. (2011). Regulating mitochondrial outer membrane proteins by ubiquitination and proteasomal degradation. Curr. Opin. Cell Biol. 23, 476–482.

Khaminets, A., Heinrich, T., Mari, M., Grumati, P., Huebner, A.K., Akutsu, M., Liebmann, L., Stolz, A., Nietzsche, S., Koch, N., et al. (2015). Regulation of endoplasmic reticulum turnover by selective autophagy. Nature 522, 354–358.

Kim, J., Kundu, M., Viollet, B., and Guan, K.-L. (2011). AMPK and mTOR regulate autophagy through direct phosphorylation of Ulk1. Nat. Cell Biol. 13, 132–141.

Komatsu, M., Chiba, T., Tatsumi, K., Iemura, S., Tanida, I., Okazaki, N., Ueno, T., Kominami, E., Natsume, T., and Tanaka, K. (2004). A novel protein-conjugating system for Ufm1, a ubiquitin-fold modifier. EMBO J. 23, 1977–1986.

Kurth, I., Pamminger, T., Hennings, J.C., Soehendra, D., Huebner, A.K., Rotthier, A., Baets, J., Senderek, J., Topaloglu, H., Farrell, S.A., et al. (2009). Mutations in FAM134B, encoding a newly identified Golgi protein, cause severe sensory and autonomic neuropathy. Nat. Genet. 41, 1179–1181.

Lake, N.J., Compton, A.G., Rahman, S., and Thorburn, D.R. (2016). Leigh syndrome: One disorder, more than 75 monogenic causes. Ann. Neurol. 79, 190–203.

Lemaire, K., Moura, R.F., Granvik, M., Igoillo-Esteve, M., Hohmeier, H.E., Hendrickx, N., Newgard, C.B., Waelkens, E., Cnop, M., and Schuit, F. (2011). Ubiquitin fold modifier 1 (UFM1) and its target UFBP1 protect pancreatic beta cells from ER stress-induced apoptosis. PLoS ONE 6, e18517.

Léon, S., and Haguenauer-Tsapis, R. (2009). Ubiquitin ligase adaptors: regulators of ubiquitylation and endocytosis of plasma membrane proteins. Exp. Cell Res. 315, 1574–1583.

Leto, D.E., Morgens, D.W., Zhang, L., Walczak, C.P., Elias, J.E., Bassik, M.C., and Kopito, R.R. (2019). Genome-wide CRISPR Analysis Identifies Substrate-Specific Conjugation Modules in ER-Associated Degradation. Mol. Cell 73, 377–389.e11.

Li, W., Xu, H., Xiao, T., Cong, L., Love, M.I., Zhang, F., Irizarry, R.A., Liu, J.S., Brown, M., and Liu, X.S. (2014). MAGeCK enables robust identification of essential genes from genome-scale CRISPR/Cas9 knockout screens. Genome Biol. 15, 554.

Li, W., Köster, J., Xu, H., Chen, C.-H., Xiao, T., Liu, J.S., Brown, M., and Liu, X.S. (2015). Quality control, modeling, and visualization of CRISPR screens with MAGeCK-VISPR. Genome Biol. 16, 281.

Liang, J.R., Lingeman, E., Ahmed, S., and Corn, J.E. (2018). Atlastins remodel the endoplasmic reticulum for selective autophagy. J. Cell Biol. 217, 3354–3367.

Lingeman, E., Jeans, C., and Corn, J.E. (2017). Production of purified casrnps for efficacious genome editing. Curr. Protoc. Mol. Biol. 120, 31.10.1-31.10.19.

Liu, J., Wang, Y., Song, L., Zeng, L., Yi, W., Liu, T., Chen, H., Wang, M., Ju, Z., and Cong, Y.-S. (2017). A critical role of DDRGK1 in endoplasmic reticulum homoeostasis via regulation of IRE1α stability. Nat. Commun. 8, 14186.

Lombardi, A.A., and Elrod, J.W. (2017). Mediating ER-mitochondrial cross-talk. Science 358, 591–592.

Mader, B.J., Pivtoraiko, V.N., Flippo, H.M., Klocke, B.J., Roth, K.A., Mangieri, L.R., and Shacka, J.J. (2012). Rotenone inhibits autophagic flux prior to inducing cell death. ACS Chem. Neurosci. 3, 1063– 1072.

Mignon-Ravix, C., Milh, M., Kaiser, C.S., Daniel, J., Riccardi, F., Cacciagli, P., Nagara, M., Busa, T., Liebau, E., and Villard, L. (2018). Abnormal function of the UBA5 protein in a case of early developmental and epileptic encephalopathy with suppression-burst. Hum. Mutat. 39, 934–938.

Milev, M.P., Grout, M.E., Saint-Dic, D., Cheng, Y.-H.H., Glass, I.A., Hale, C.J., Hanna, D.S., Dorschner, M.O., Prematilake, K., Shaag, A., et al. (2017). Mutations in TRAPPC12 manifest in progressive childhood encephalopathy and golgi dysfunction. Am. J. Hum. Genet. 101, 291–299.

Nahorski, M.S., Maddirevula, S., Ishimura, R., Alsahli, S., Brady, A.F., Begemann, A., Mizushima, T., Guzmán-Vega, F.J., Obata, M., Ichimura, Y., et al. (2018). Biallelic UFM1 and UFC1 mutations expand the essential role of ufmylation in brain development. Brain 141, 1934–1945.

Nguyen, T.N., Padman, B.S., and Lazarou, M. (2016). Deciphering the molecular signals of pink1/parkin mitophagy. Trends Cell Biol. 26, 733–744.

Nordgren, M., Francisco, T., Lismont, C., Hennebel, L., Brees, C., Wang, B., Van Veldhoven, P.P., Azevedo, J.E., and Fransen, M. (2015). Export-deficient monoubiquitinated PEX5 triggers peroxisome removal in SV40 large T antigen-transformed mouse embryonic fibroblasts. Autophagy 11, 1326–1340.

Novarino, G., Fenstermaker, A.G., Zaki, M.S., Hofree, M., Silhavy, J.L., Heiberg, A.D., Abdellateef, M., Rosti, B., Scott, E., Mansour, L., et al. (2014). Exome sequencing links corticospinal motor neuron disease to common neurodegenerative disorders. Science 343, 506–511.

Phillips, A.R., Suttangkakul, A., and Vierstra, R.D. (2008). The ATG12-conjugating enzyme ATG10 Is essential for autophagic vesicle formation in Arabidopsis thaliana. Genetics 178, 1339–1353.

Pickrell, A.M., and Youle, R.J. (2015). The roles of PINK1, parkin, and mitochondrial fidelity in Parkinson’s disease. Neuron 85, 257–273.

Pilsl, A., and Winklhofer, K.F. (2012). Parkin, PINK1 and mitochondrial integrity: emerging concepts of mitochondrial dysfunction in Parkinson’s disease. Acta Neuropathol. 123, 173–188.

Rismanchi, N., Soderblom, C., Stadler, J., Zhu, P.-P., and Blackstone, C. (2008). Atlastin GTPases are required for Golgi apparatus and ER morphogenesis. Hum. Mol. Genet. 17, 1591–1604.

Ruggiano, A., Foresti, O., and Carvalho, P. (2014). ER-associated degradation: protein quality control and beyond. J. Cell Biol. 204, 869–879.

Schuck, S., Prinz, W.A., Thorn, K.S., Voss, C., and Walter, P. (2009). Membrane expansion alleviates endoplasmic reticulum stress independently of the unfolded protein response. J. Cell Biol. 187, 525–536.

Schwarz, D.S., and Blower, M.D. (2016). The endoplasmic reticulum: structure, function and response to cellular signaling. Cell. Mol. Life Sci. 73, 79–94.

Smith, M.D., Harley, M.E., Kemp, A.J., Wills, J., Lee, M., Arends, M., von Kriegsheim, A., Behrends, C., and Wilkinson, S. (2018). CCPG1 Is a Non-canonical Autophagy Cargo Receptor Essential for ER-Phagy and Pancreatic ER Proteostasis. Dev. Cell 44, 217–232.e11.

Szklarczyk, D., Franceschini, A., Wyder, S., Forslund, K., Heller, D., Huerta-Cepas, J., Simonovic, M., Roth, A., Santos, A., Tsafou, K.P., et al. (2015). STRING v10: protein-protein interaction networks, integrated over the tree of life. Nucleic Acids Res. 43, D447–52.

Taanman, J.-W. (1999). The mitochondrial genome: structure, transcription, translation and replication. Biochimica et Biophysica Acta (BBA) – Bioenergetics 1410, 103–123.

Toyama, E.Q., Herzig, S., Courchet, J., Lewis, T.L., Losón, O.C., Hellberg, K., Young, N.P., Chen, H., Polleux, F., Chan, D.C., et al. (2016). Metabolism. AMP-activated protein kinase mediates mitochondrial fission in response to energy stress. Science 351, 275–281.

Walczak, C.P., Leto, D.E., Zhang, L., Riepe, C., Muller, R.Y., DaRosa, P.A., Ingolia, N.T., Elias, J.E., and Kopito, R.R. (2019). Ribosomal protein RPL26 is the principal target of UFMylation. Proc Natl Acad Sci USA.

Wang, S., Tukachinsky, H., Romano, F.B., and Rapoport, T.A. (2016). Cooperation of the ER-shaping proteins atlastin, lunapark, and reticulons to generate a tubular membrane network. Elife 5.

Wang, X., Winter, D., Ashrafi, G., Schlehe, J., Wong, Y.L., Selkoe, D., Rice, S., Steen, J., LaVoie, M.J., and Schwarz, T.L. (2011). PINK1 and Parkin target Miro for phosphorylation and degradation to arrest mitochondrial motility. Cell 147, 893–906.

Wartosch, L., Günesdogan, U., Graham, S.C., and Luzio, J.P. (2015). Recruitment of VPS33A to HOPS by VPS16 Is Required for Lysosome Fusion with Endosomes and Autophagosomes. Traffic 16, 727–742.

Wei, Y., and Xu, X. (2016). Ufmylation: A unique & fashionable modification for life. Genomics Proteomics Bioinformatics 14, 140–146.

Wu, J., Lei, G., Mei, M., Tang, Y., and Li, H. (2010). A novel C53/LZAP-interacting protein regulates stability of C53/LZAP and DDRGK domain-containing Protein 1 (DDRGK1) and modulates NF-kappaB signaling. J. Biol. Chem. 285, 15126–15136.

Xiao, B., Deng, X., Zhou, W., and Tan, E.-K. (2016). Flow Cytometry-Based Assessment of Mitophagy Using MitoTracker. Front. Cell. Neurosci. 10, 76.

Yoo, H.M., Kang, S.H., Kim, J.Y., Lee, J.E., Seong, M.W., Lee, S.W., Ka, S.H., Sou, Y.-S., Komatsu, M., Tanaka, K., et al. (2014). Modification of ASC1 by UFM1 is crucial for ERα transactivation and breast cancer development. Mol. Cell 56, 261–274.

Youle, R.J., and Narendra, D.P. (2011). Mechanisms of mitophagy. Nat. Rev. Mol. Cell Biol. 12, 9–14.

Yuan, H.-X., Russell, R.C., and Guan, K.-L. (2013). Regulation of PIK3C3/VPS34 complexes by MTOR in nutrient stress-induced autophagy. Autophagy 9, 1983–1995.

Zhang, J., Tripathi, D.N., Jing, J., Alexander, A., Kim, J., Powell, R.T., Dere, R., Tait-Mulder, J., Lee, J.-H., Paull, T.T., et al. (2015). ATM functions at the peroxisome to induce pexophagy in response to ROS. Nat. Cell Biol. 17, 1259–1269.

Zhao, G., Zhu, P.-P., Renvoisé, B., Maldonado-Báez, L., Park, S.H., and Blackstone, C. (2016). Mammalian knock out cells reveal prominent roles for atlastin GTPases in ER network morphology. Exp. Cell Res. 349, 32–44.

